# Quantitative modeling of SARS-CoV-2 replication reveals phase-specific bottlenecks and antiviral targets

**DOI:** 10.64898/2026.06.23.733955

**Authors:** Simon T. Herrmann, Timon Kapischke, Saskia Westhoven, Natalie Heinen, Luca D. Bertzbach, Toni L. Meister, Barbara Sitek, Thilo Bracht, Stephanie Pfaender, Lars Kaderali

## Abstract

SARS-CoV-2 replication depends on a tightly coordinated series of intracellular processes that remain incompletely quantified. Here, we integrated high-resolution time-resolved measurements of viral RNA, protein expression, and infectious virion production with mechanistic mathematical modeling to obtain a quantitative description of the viral replication cycle in human lung cells. Using transcriptomic, proteomic, and infectivity data collected over the first 24 hours of infection, we calibrated an ordinary differential equation model that captures genomic and subgenomic RNA synthesis, viral protein production, virion assembly, and virus release. The model accurately reproduced the observed replication dynamics and enabled estimation of kinetic parameters that are difficult to measure experimentally. Sensitivity analysis identified viral RNA replication and non-structural protein maturation as dominant determinants of viral replication efficiency. To assess predictive power, the model was challenged with independent antiviral perturbation experiments using remdesivir, nirmatrelvir, and montelukast. Model predictions closely matched experimentally observed treatment responses and correctly reproduced drug interaction effects during combination therapy. Furthermore, comparison of alternative mechanistic hypotheses supported NSP5 rather than NSP1 as the primary antiviral target of montelukast. Together, these results establish a predictive framework for dissecting intracellular coronavirus replication and evaluating antiviral intervention strategies.

## Introduction

Severe acute respiratory syndrome coronavirus 2 (SARS-CoV-2), the causative agent of COVID-19, triggered a global pandemic beginning in late 2019 and has since resulted in hundreds of millions of confirmed infections and millions of deaths worldwide. Beyond its immediate public health impact, the virus has become a focal point for studying the molecular mechanisms governing coronavirus replication and antiviral intervention strategies. While vaccination remains the most effective means in preventing severe disease, understanding the intracellular replication cycle of SARS-CoV-2 is critical for identifying rate-limiting processes and regulatory bottlenecks that can be exploited by antiviral interventions [1]. SARS-CoV-2 is an enveloped positive-sense single-stranded RNA virus. It has the largest genome among all human pathogenic RNA viruses, with a length of approximately 30 kb. Although it primarily infects epithelial cells of the respiratory tract [2, 3], multi-organ tropism has been reported, including involvement of the heart, kidneys, intestines and, to a lesser extent, the central nervous system [4, 5]. Coronaviruses (CoVs) are characterized by discontinuous transcription, which generates a nested set of subgenomic RNAs (sgRNAs) encoding structural and accessory proteins. SARS-CoV-2 encodes four structural proteins: nucleocapsid (N), spike (S), membrane (M), and envelope (E); seven accessory proteins (ORF3a, ORF3c, ORF6, ORF7a, ORF7b, ORF8, and ORF9b); and two polyproteins (pp1a and pp1ab) that are further proteolytically cleaved into 16 non-structural proteins (NSP1 through NSP16) [6] (S1 Fig.).

Viral entry is initiated by binding of the spike protein to the angiotensin-converting enzyme 2 (ACE2) receptor on the surface of susceptible host cells [3], followed by proteolytic activation of the spike protein by host proteases such as transmembrane serine protease 2 (TMPRSS2) or endosomal cathepsins, enabling membrane fusion and release of the viral genome into the host cell cytoplasm [4, 7]. Following entry, the positive-sense, single stranded genomic RNA (gRNA) is uncoated and released into the cytoplasm. The 5’-proximal region of the gRNA, corresponding to ORF1a/1ab, is translated into two large polyproteins, pp1a and pp1ab, which are cleaved by the viral papain-like protease (within NSP3) and the main protease NSP5 into the 16 NSPs (S1 Fig.). Several of these NSPs, including NSP3, NSP4 and NSP6, drive the formation of double-membrane vesicles (DMVs) derived from the endoplasmic reticulum (ER), which serve as the central site of viral RNA synthesis [8]. Within these compartments, SARS-CoV-2 employs a distinctive discontinuous transcription mechanism that gives rise to a nested set of sgRNAs in addition to full-length gRNA. Both gRNA and sgRNAs are translocated to the cytoplasm, where the gene located at the 5’ end of each sgRNA is translated into structural or accessory proteins. While accessory proteins contribute to immune evasion and manipulation of host cell processes, structural proteins are incorporated into newly assembled virions together with gRNA. Assembly occurs in the ER-Golgi intermediate compartment, where N-coated gRNA interacts with membrane-associated M, E, and S proteins. Mature virions are subsequently released through vesicular trafficking pathways, including exocytic and lysosome-derived routes, enabling infection of neighboring cells and continuation of the replication cycle [6]. While the individual stages of the SARS-CoV-2 replication cycle have been characterized in considerable detail, their quantitative interplay over time remains difficult to assess experimentally. Mathematical modeling provides a framework for integrating these processes and analyzing their contribution to viral replication dynamics.

The COVID-19 pandemic highlighted the need for a deeper mechanistic understanding of CoV biology in order to improve preparedness for future outbreaks and emerging viral threats. In this context, the combination of quantitative measurements and dynamic mathematical modeling has provided important insights into viral infection processes [5]. The increasing availability of quantitative, time-resolved intracellular data offers an opportunity to investigate these mechanisms systematically. In particular, recent advances in transcriptomics, proteomics, and quantitative virology enable simultaneous observation of multiple stages of the viral life cycle, creating new opportunities for data-driven mechanistic modeling. Given the complexity of the viral replication cycle, accurately capturing intracellular dynamics requires both rich, time-resolved experimental data and mechanistic structure. Ordinary differential equation (ODE)-based models, in particular, provide a natural framework for translating experimental observations into quantitative estimates of reaction rates, feedback mechanisms, and regulatory dependencies that are otherwise not directly observable. By explicitly representing key stages of the viral replication cycle, such models enable a systems-level understanding of replication dynamics, facilitate the mechanistic interpretation of experimental perturbations, and help identify rate-limiting processes that may serve as targets for antiviral intervention. To date, only a limited number of mathematical models describing the SARS-CoV-2 replication cycle have been published [9–15]. Although these studies have provided valuable insights, most were calibrated using relatively limited experimental datasets, potentially restricting parameter identifiability, predictive accuracy, and generalizability.

In this study, we combine time-resolved experimental measurements with mechanistic modeling to characterize the intracellular replication dynamics of SARS-CoV-2 in human lung cells. Using multi-omics data generated from infected A549 cultures, overexpressing ACE2 and TMPRSS2 (A/T), together with measurements of infectious viral titers, we developed a comprehensive ordinary differential equation (ODE) model describing the coordinated production of viral RNA species, proteins, and infectious virions. Calibration of the model against experimental observations yields a quantitative representation of the viral replication cycle and enables estimation of key kinetic parameters that are not directly accessible experimentally. To assess the predictive power of the model, we evaluated its performance under independent antiviral perturbations targeting distinct stages of the replication cycle. Specifically, we investigated three FDA-approved compounds: nirmatrelvir, remdesivir and montelukast [16]. The antiviral mechanisms of nirmatrelvir and remdesivir are well established, whereas the antiviral mechanism for montelukast remains incompletely understood [16–18]. By challenging the model with compounds acting at different points in the viral replication cycle, we assess its ability to generalize beyond the conditions used for parameter estimation and to reproduce independent experimental observations. Together, this integrative framework provides a quantitative description of SARS-CoV-2 replication and a foundation for evaluating antiviral interventions, identifying rate-limiting processes, and generating experimentally testable hypotheses regarding CoV biology.

## Results

### Construction of a mechanistic ODE model capturing the intracellular SARS-CoV-2 replication dynamics

We developed a mechanistic ODE model describing the intracellular replication cycle of SARS-CoV-2. Building on our previous work on hepatitis C virus and dengue virus [19, 20], the model was extended to incorporate biological features specific to CoVs, including discontinuous transcription and the production of sgRNAs. Compared with previously published intracellular SARS-CoV-2 replication models [9–15], our approach was calibrated and validated using a comprehensive 24 h time-resolved multi-omics dataset. The model explicitly represents genomic and sgRNA species, viral proteins, intracellular, as well as released infectious particles, thereby capturing the major stages of the viral replication cycle from genome replication to virion egress. Based on mass action kinetics, the model comprises 59 ODEs and 49 parameters describing RNA synthesis, protein translation, virion assembly and virus release. A schematic overview of the model structure is shown in Fig. 1.

**Figure 1:**
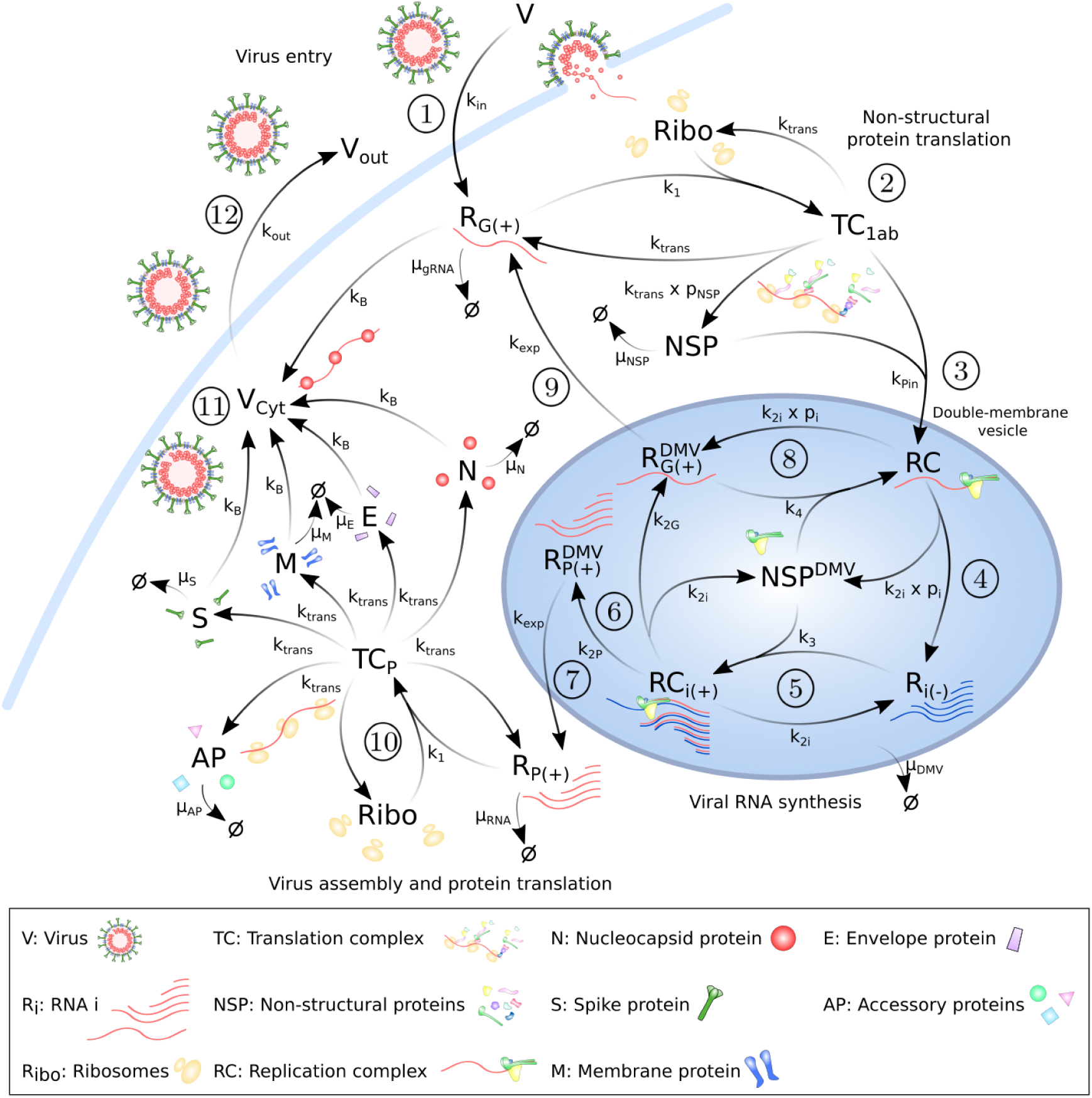
Schematic overview of the SARS-CoV-2 ODE replication model. ① The virus (*V*) binds to the ACE2 receptor and enters the cell at a rate *k_in_*, releasing positive-sense genomic RNA *R_G_*_(+)_ into the cytoplasm. ② Ribosomes *Ribo* bind to *R_G_*_(+)_ at rate *k*_1_, forming the translation complex *TC*_1_*_ab_*, which translates non-structural proteins *NSP* at rate *k_trans_* with probability *p_NSP_* of generating functional NSPs. *NSP* are degraded at a rate *µ_NSP_*. ③ *NSP* and *TC*_1_*_ab_* are imported into double-membrane vesicles (DMVs), where they form the replication complex *RC*. ④ Within *RC*, *R_G_*_(+)_ serves as template for the synthesis of negative-strand RNA *R_i_*_(_*_−_*_)_ at rate *k*_2_*_i_*. Transcription is directed towards genomic or subgenomic RNA synthesis according to probabilities *p_i_*. The *RC* subsequently disassembles into 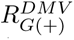 and non-structural proteins *NSP^DMV^*. ⑤ Negative-strand RNAs initiate formation of transcription-replication complexes *RC_i_*_(+)_ at rate *k*_3_, which synthesizes positive-strand RNAs at rate *k*_2_*_i_*. ⑥ They synthesize new plus-strand gRNAs 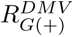 and sgRNAs 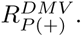 ⑦ Positive sgRNAs are exported to the cytoplasm at rate *k_exp_*. ⑧ Newly synthesized genomic RNAs either re-enter the replication cycle or ⑨ are exported to the cytoplasm at rate *k_exp_*. All DMV-associated species are degraded at rate *µ_DMV_*. ⑩ Ribosomes *Ribo* bind (*k*_1_) positive strand sgRNAs *R_P_* _(+)_ and form a translation initiation complex *TC_P_*, which translate (*k_trans_*) structural proteins *SP ∈ {N, S, M, E}* and accessory proteins *AP ∈ {*3*a,* 6, 7*a,* 8*}*, which are degraded at rates *µ_SP_* or *µ_AP_*, respectively. Only accessory proteins for which quantitative measurements were available were explicitly represented in the model. ⑪ Structural proteins and genomic RNA assemble into virions at rate *k_B_*, which are then ⑫ exported from the cell at rate *k_out_*. The index *i ∈ {G, N, E, M, S,* 3*a,* 6, 7*a,* 8*}* denotes genomic RNA and the individual subgenomic RNA species represented in the model, corresponding to the proteins *P ∈ {N, E, M, S,* 3*a,* 6, 7*a,* 8*}*.

### Multi-omics time-series measurements characterize viral replication dynamics

To characterize SARS-CoV-2 replication dynamics at high temporal resolution, we performed a dense time-course experiment in infected A549 (A/T) cells over the first 24 hours post infection (hpi). To obtain quantitative measurements across multiple stages of the viral replication cycle, we combined transcriptomic, proteomic, and infectivity measurements from the same infection experiment. We collected samples hourly during the first 8 hpi and every two hours thereafter until 24 hpi (Fig. 2). Viral transcriptome dynamics were quantified by RT-qPCR measurements of gRNA and E sgRNA copy numbers, complemented by sequencing-based quantification of additional sgRNA species (Fig. 2a, 2b and S2 Fig.). Viral protein abundance was quantified by liquid chromatography-tandem mass spectrometry (LC-MS/MS), with absolute nucleocapsid levels determined using calibrated western blot analyses (Fig. 2c and S3 Fig.). Finally, intracellular and extracellular infectious virions were quantified by endpoint dilution assays (TCID_50_/mL), enabling temporal characterization of virus production and release (Fig. 2d and 2e).

**Figure 2:**
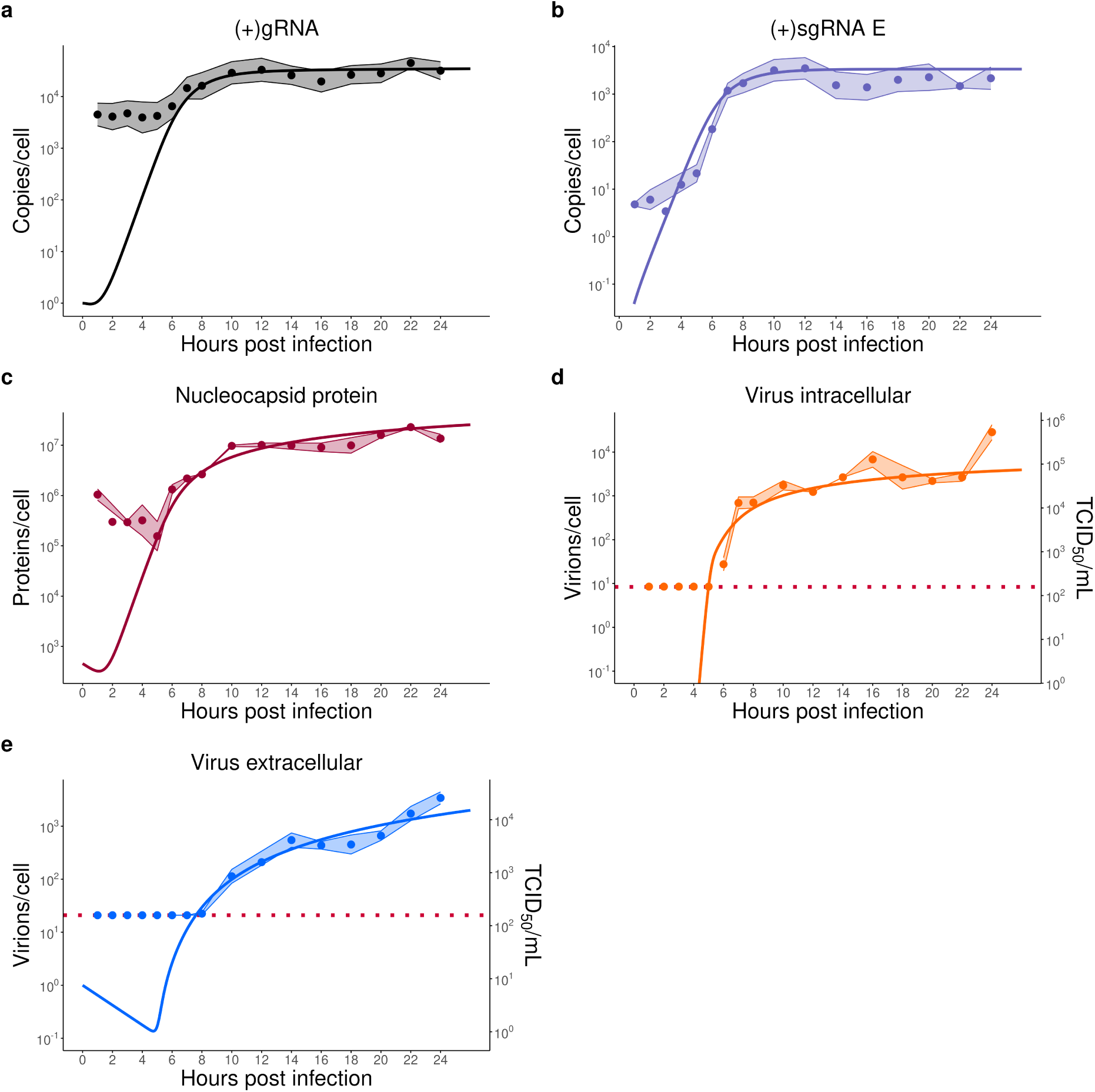
Experimental datasets and model fit. In the 24 hour kinetic experiment, samples were harvested at the indicated time points and analyzed. Additional measurements are presented in Supplementary Figs. S2 and S3. Mean experimental values are shown as dots, and shaded areas indicate the 95% confidence interval. The solid lines represent the fitted model trajectories obtained after parameter estimation. **a** and **b** Viral RNA levels quantified by RT-qPCR. gRNA and sgRNA E were measured over time and converted to copies per cell based on normalization to GAPDH. **c** Viral protein expression over time was quantified by western blotting. Nucleocapsid protein abundance was quantified relative to *β*-actin expression and calibrated using a nucleocapsid protein dilution series as a standard curve. **d** and **e** Infectivity titration (TCID_50_/mL) quantifying infectious virus particles during the first 24 hpi, shown separately for intracellular **d** and extracellular **e** fractions. The left y-axis shows the model-predicted number of infectious virions per cell (orange and blue lines), while the right y-axis shows corresponding TCID_50_/mL values measured from experimental samples. Dotted red lines indicate the detection limit of the TCID_50_ assay. Data are derived from n=5 independent experiments.

Our transcriptomic dynamics revealed a biphasic pattern. We detected gRNA levels throughout the entire time course, showing an initially stable level up to approximately 5 hpi, followed by a pronounced increase, which reached a plateau at around 10 hpi, after which levels remained relatively constant until the end of the observation period (Fig. 2a). sgRNA species were initially near or below the limit of detection until approximately 5 hpi, after which they increased exponentially, reaching a plateau around 12 hpi and subsequently stabilizing (Fig. 2b). Notably, this temporal pattern was consistent across all measured sgRNA species, suggesting highly co-ordinated expression kinetics across viral transcripts. Proteomic measurements followed similar kinetics, with an apparent delay of approximately one hour relative to sgRNA accumulation. At 6 hpi, intracellular infectious viral particles became detectable by end-point dilution assay. Viral particle formation increased over the following hour, indicating exponential growth, before reaching a plateau or increasing only marginally until the end of the sampling period. In contrast, we only detected infectious viral particles released into the supernatant at 10 hpi, indicating an approximately two hour lag between the first detection of intracellular infectious virions and their appearance in the extracellular compartment (Fig. 2d and 2e). By integrating these multi-omics trajectories of transcriptomics and proteomics combined with infectious particle measurements, we were able to fit our mathematical model (Fig. 1) that quantitatively describes the SARS-CoV-2 replication cycle representing the average dynamics of a single infected cell. Model calibration further enabled estimation of quantities not directly observable experimentally. The fitted model predicts that an average infected cell produces 5321 infectious virions within 24 hpi, of which approximately 31% (1655 virions) are released into the extracellular compartment. The first particle is released at approximately 5.5 hpi (for additional kinetics parameters, see S1 Table). This high-resolution time course enabled precise parameterization of the model, capturing key kinetic features such as early transcriptional activation, exponential growth amplification phases, viral assembly, and delayed release dynamics. Collectively, these data provide a quantitative description of the temporal progression of SARS-CoV-2 replication revealing a cascade of temporally separated events characterized by increases in gRNA and sgRNA at approximately 5 hpi, viral protein accumulation and intracellular virion production at approximately 6 hpi, and extracellular virion release at approximately 10 hpi. This sequential progression establishes the biological foundation of the entire model and provides a rich experimental framework for mechanistic model calibration and validation.

### Sensitivity analyses identify dominant kinetic processes shaping viral replication output

Global sensitivity analysis using the extended Fourier amplitude sensitivity test (eFAST; Fig. 3) revealed that viral replication dynamics are dominated by a small subset of highly influential parameters, reflecting key bottlenecks in viral replication efficiency. Across all analyzed time points, translation rate (*k_trans_*), replication rate (*k_RdRp_*) and initial ribosome abundance (*Ribo_init_*) consistently ranked among the most influential parameters, indicating that protein synthesis capacity and RNA replication are major determinants of viral replication efficiency. In addition, the degradation rate within DMVs (*µ_DMV_*) and the gRNA degradation (*µ_gRNA_*) strongly affected intracellular accumulation of positive-sense genomic RNA. For virion assembly and release, the egress of virions (*k_out_*) and the number of structural membrane proteins per virion (*n_M_*) emerged as the most important parameters. Our time-resolved analysis further highlighted the importance of the early-stage parameters *p_NSP_*and *k*_1_ governing early NSP production. The parameter *k*_1_, describing the binding of genomic RNA to ribosomes and thereby the initiation of ORF1a/1ab translation, exerted its strongest influence during the early phase of infection. Similiarly, *p_NSP_* representing the probability of generating functional NSPs via NSP5-induced autoproteolysis. The decreasing influence of both parameters at later time points, indicating that early NSP production is particularly important during the establishment of infection, whereas later stages become increasingly governed by downstream replication processes. Together, these analyses indicate that SARS-CoV-2 replication is governed by a limited number of dominant processes, particularly RNA replication, protein synthesis, and virion production, which represent potential points of vulnerability within the viral replication cycle. Notably, two of the most influential parameters identified by the sensitivity analysis, *k_RdRp_* and *p_NSP_*, correspond to the targets of remdesivir and nirmatrelvir, respectively, enabling direct experimental evaluation of the model predictions.

**Figure 3:**
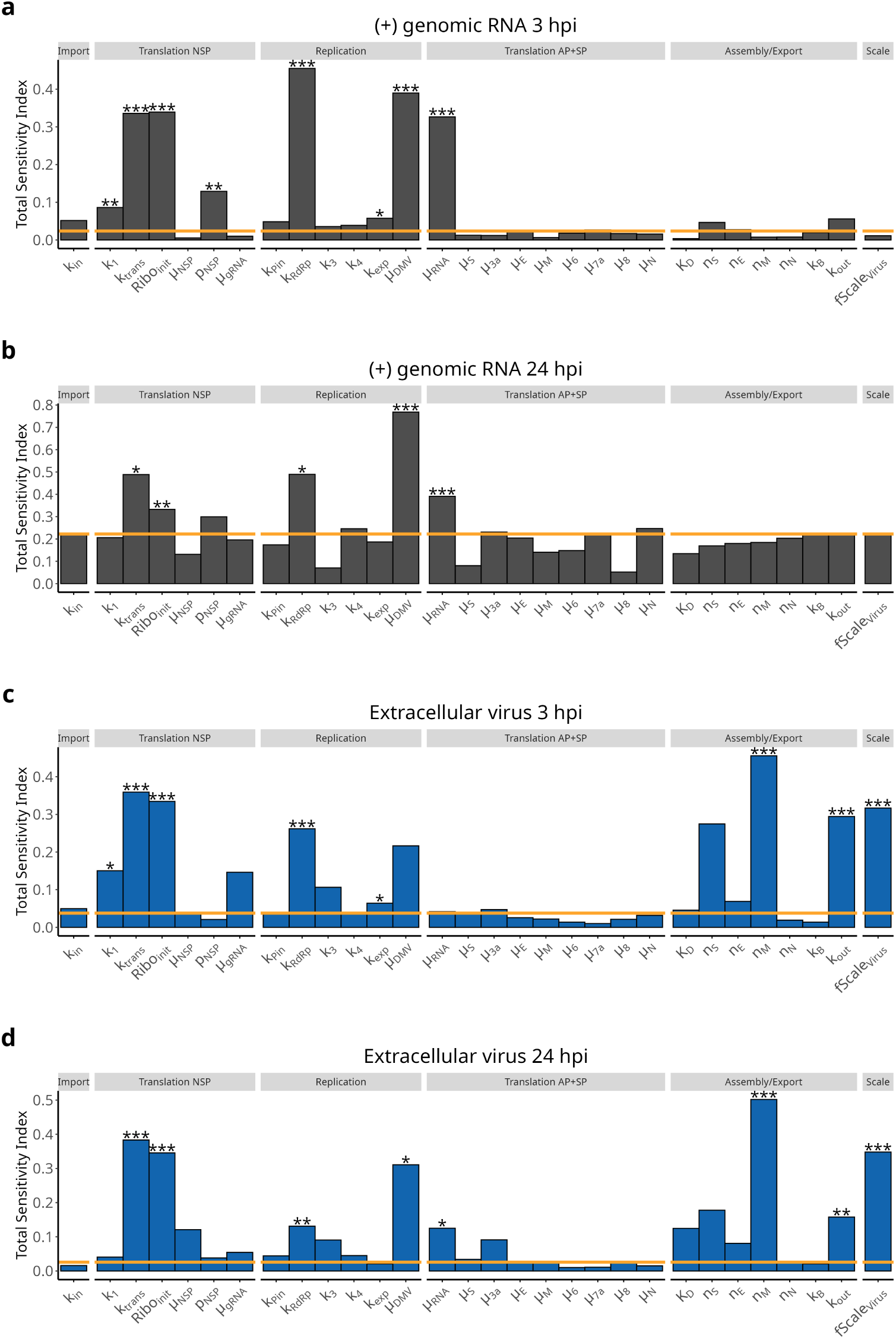
Global sensitivity analysis of model outputs. Sensitivity analysis was performed using the extended Fourier Amplitude Sensitivity Test (eFAST) at 3 and 24 hpi. Parameters are grouped by functional category: import, NSP translation, replication, accessory and structural protein translation, assembly/export, and scaling. Each bar represents the total sensitivity index. The orange line indicates the sensitivity index obtained for a dummy parameter not included in the model. Parameters with sensitivity indices at or below this level were considered indistinguishable from background sensitivity. In all panels, statistical significance was assessed by comparing parameter sensitivities against the dummy parameter using a t-test (p-values: *** *≤* 0.001, ** *≤* 0.01, * *≤* 0.05). **a**, **b** Total intracellular positive-sense genomic RNA ((+)gRNA). Total sensitivity indices describing the influence of individual parameters on intracellular (+)gRNA accumulation at 3 and 24 hpi. **c**, **d** Extracellular infectious virions. Total sensitivity indices describing the influence of individual parameters on extracellular virus production at 3 and 24 hpi.

### Model accurately predicts viral replication under targeted antiviral inhibition

To evaluate the predictive capability of the model and to assess whether the inferred mechanisms generalize beyond the calibration dataset, we challenged the model with antiviral perturbations targeting defined stages of the replication cycle. Dose-response experiments were performed to determine compound-specific IC_50_ values for remdesivir, montelukast, and nirmatrelvir (Fig. 4a–c). Drug treatment was incorporated into the model using a Hill-type inhibition function that modulated the corresponding target parameters. Remdesivir was modeled as inhibition of RNA replication (*k_RdRp_*), nirmatrelvir as inhibition of NSP maturation (*p_NSP_*), and montelukast as inhibition of either NSP1-mediated translation initiation (*k*_1_) or NSP5-dependent NSP processing (*p_NSP_*). Experimental time-course measurements of extracellular infectivity, representing de-novo formed SARS-CoV-2 particles released into the supernatant under drug treatment, were compared with model-based inhibition predictions (Fig. 4d–f). Model predictions closely reproduced the experimentally observed kinetics of extracellular virus production across all treatment conditions. Similar agreement was observed for intracellular infectivity measurements. Together, these results indicate that the model can accurately reproduce perturbed conditions not used during initial parameterization. Because the antiviral mechanism of montelukast remains uncertain, we compared simulations assuming inhibition of either NSP1 or NSP5. The NSP5-targeting model reproduced the observed kinetics more accurately, particularly during the early phase of infection (S4 Fig.), and yielded lower prediction errors (S2 Table), supporting NSP5 as the more likely antiviral target. From the most sensitive parameters identified in the sensitivity analysis (Fig. 3), only two were inhibited: *p_NSP_*and *k_RdRp_*. However, *p_NSP_* significantly affects the replication cycle only during the first few hours, which is reflected in the simulated efficacy. Consistent with the sensitivity analysis, inhibition of *k_RdRp_*produced a larger reduction in viral output than inhibition of *p_NSP_*, whose influence is largely restricted to the early stages of infection. For remdesivir monotherapy simulations to investigate the discrepancy between nominal IC_50_-based simulations and the observed kinetics, we estimated the effective intracellular inhibition required to reproduce the experimental data (S5 Fig.). An effective concentration of 91.56 nM yielded the best agreement, corresponding to approximately 37.5% inhibition of *k_RdRp_*. We also examined how the drugs affect the dynamics of viral replication over time. The model predicts that none of the drugs permanently reduce viral load; instead, they only delay the timing of the viral peak (S6 Fig.). This suggests that drug treatment alone may be insufficient to control the infection. Together, these results demonstrate that the model accurately predicts viral replication dynamics under pharmacological perturbation and can discriminate between competing mechanistic hypotheses regarding antiviral mode of action.

**Figure 4:**
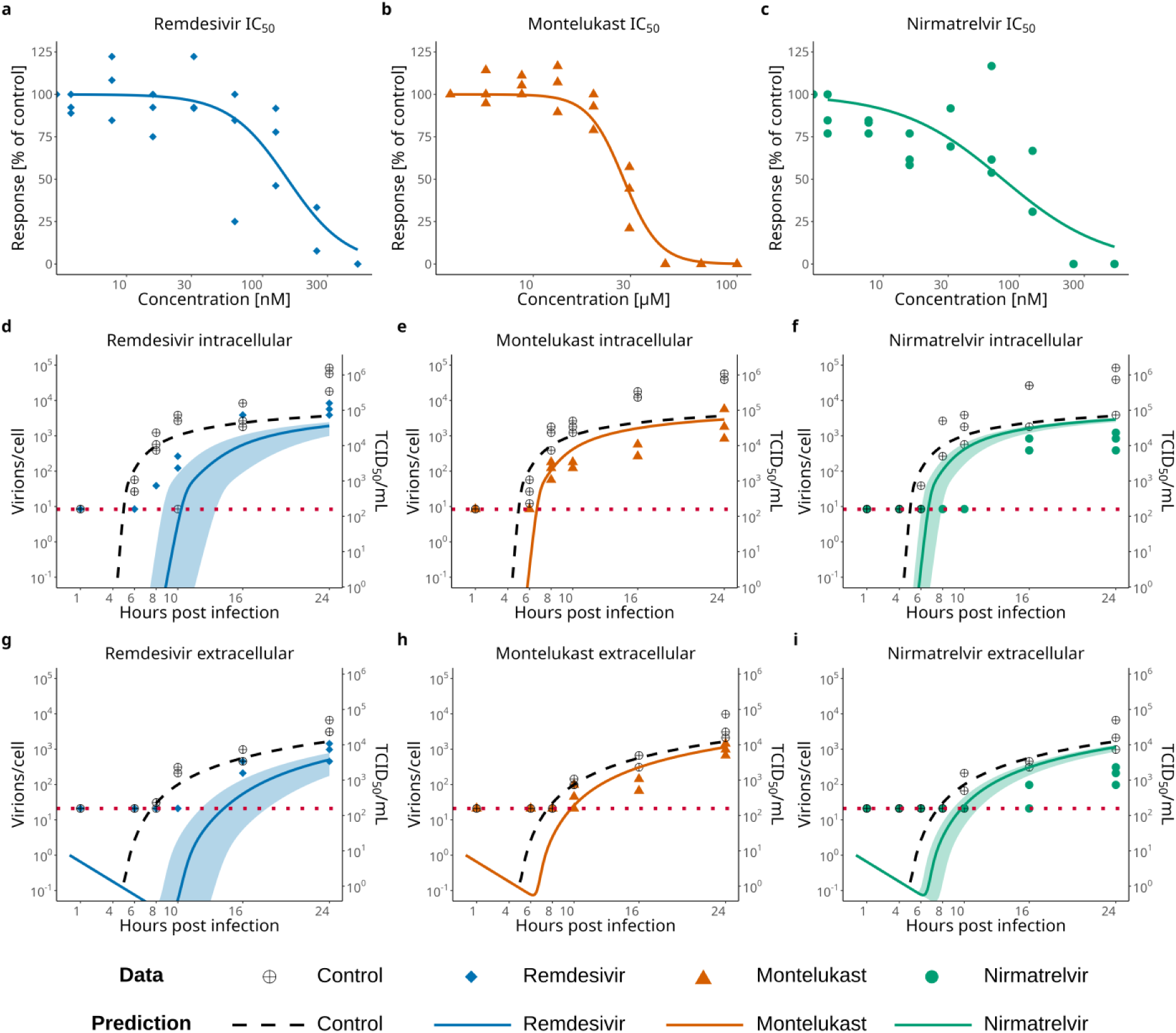
Experimental measurements and model predictions of drug treatment dynamics. **a** to **c**: Dose-response experiments used to determine compound-specific IC_50_ values for **a** remdesivir (IC_50_=152.6 nM), **b** montelukast (IC_50_=27.94 µM), and **c** nirmatrelvir (IC_50_=75.56 nM) against SARS-CoV-2 replication in infected A549 (A/T) cells. Compound concentrations are shown on the x-axis, and the y-axis indicates viral replication as a percentage relative to vehicle controls. Experimental data points represent three independent experiments, and the solid line indicates the four-parameter logistic fit. These experimentally determined IC_50_ values were subsequently used to parameterize the antiviral inhibition functions in the model. **d** to **f** : Kinetic experiments assessing extracellular titers over time following treatment with **d** remdesivir (blue), **e** montelukast (orange), and **f** nirmatrelvir (green) at their respective IC_50_ concentrations. Experimental data points are shown (IC_50_/mL, right y-axis, dotted red lines indicate the detection limit), while the model prediction is shown as a continuous line (virions/cell, left y-axis). Vehicle controls are shown for comparison. **g** to **i**: Corresponding kinetic measurements of intracellular infectivity. As above, points represent experimental measurements and the solid line represents the model-predicted kinetics under drug treatment. Shaded areas indicate model prediction uncertainty propagated from the 95% confidence interval of the experimentally determined IC_50_ values shown in panel **a** to **c**. Model predictions were generated without re-estimation of replication parameters and were compared against independent kinetic measurements under antiviral treatment.

### Mechanistic modeling predicts combination therapy responses and identifies NSP5 as the likely target of montelukast

We next experimentally characterized pairwise drug combinations and compared the resulting synergy landscapes with model predictions generated under identical conditions. ZIP synergy scores were computed for both experimental and *in-silico* data to quantify potential synergistic, additive, and or antagonistic effects (Fig. 5a–f). The predicted synergy landscapes closely matched the *in-vitro* observations, correctly reproducing both synergistic and antagonistic interactions across all drug combinations. Because the molecular target of montelukast remains uncertain, we compared model predictions assuming inhibition of either NSP1 or NSP5. The two hypotheses produced distinct synergy patterns when combined with nirmatrelvir (S7 and S8 Figs.). Under the NSP1-targeting assumption, the model predicted a synergistic interaction, whereas inhibition of NSP5 resulted in a predominantly non-synergistic response. The latter was in close agreement with the experimental data (Fig. 5c and 5f), providing additional evidence that NSP5 is the primary antiviral target of montelukast. To investigate the molecular basis of this observation, we performed docking simulations of montelukast using the high-resolution NSP5 crystal structure 8DZ2 as a template (Fig. 5g) [21]. Interestingly, this simulation showed that the 10 highest-scoring docking poses all localized montelukast to the same binding pocket occupied by nirmatrelvir (best fit shown in Fig. 5h and 5i). This overlap suggests direct competition for the NSP5 active site and provides a plausible mechanistic explanation for the antagonistic interaction observed experimentally and predicted by the model. Published studies have reported similar binding interactions, providing additional support for the proposed binding mode [17, 21]. Together, these results demonstrate that the model can accurately predict combination therapy responses, discriminate between competing mechanistic hypotheses, and generate experimentally testable insights into antiviral drug action.

**Figure 5:**
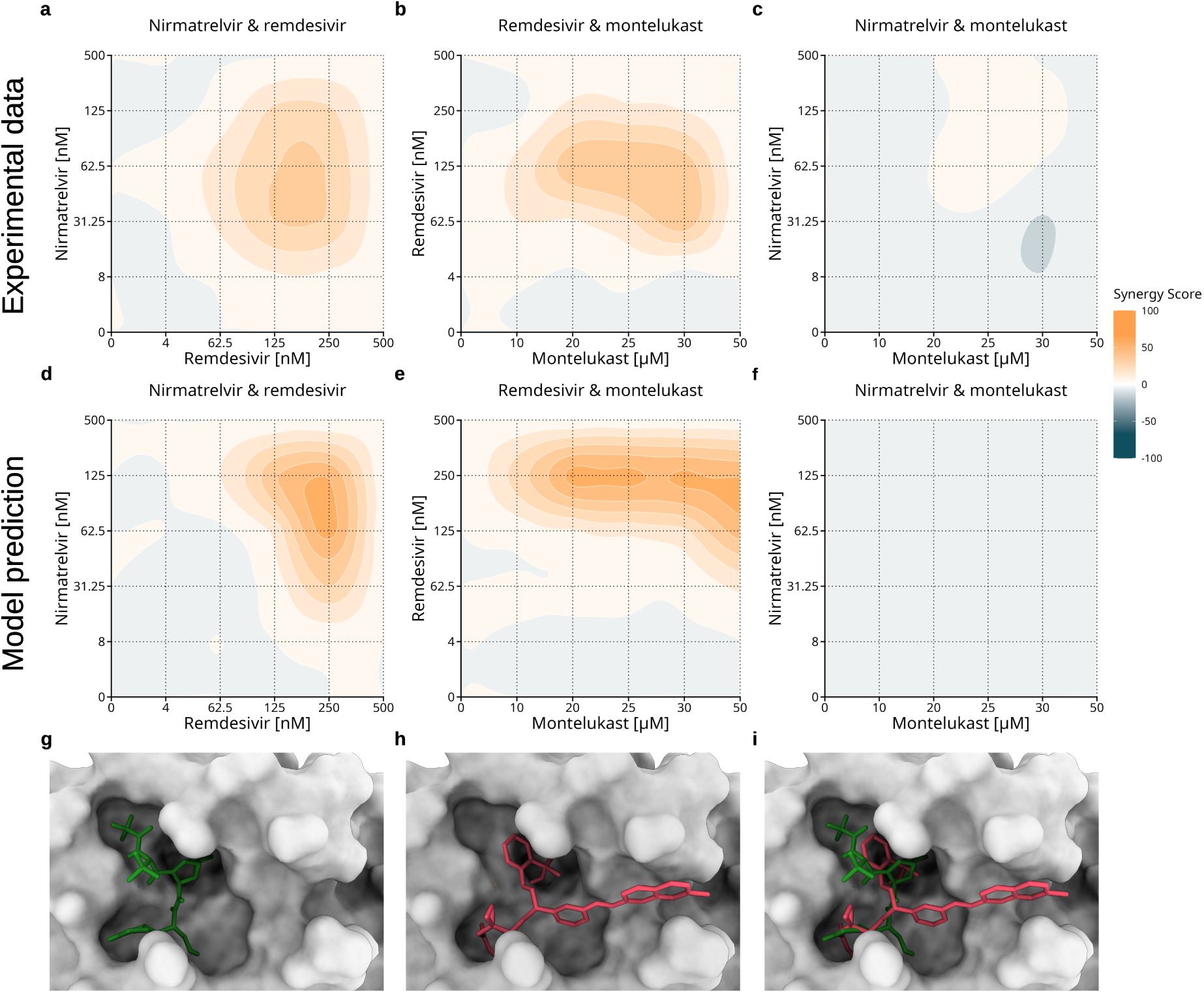
Prediction of antiviral combination effects and structural analysis of montelukast binding. **a** to **c**: Experimental synergy landscapes for the indicated drug combinations in SARS-CoV-2 infected A549 (A/T) cells. Synergy was quantified using the ZIP (zero interaction potency) model. Heatmaps show ZIP synergy scores for remdesivir and nirmatrelvir (**a**; mean ZIP = 8.75), remdesivir and montelukast (**b**; mean ZIP = 9.89), and nirmatrelvir and montelukast (**c**; mean ZIP = -1.8). Positive values (orange) indicate synergistic interactions, values near zero indicate additive effects, and negative values (blue) indicate antagonism. **d** to **f** : Model-predicted synergy landscapes. Calculations are shown for remdesivir and nirmatrelvir (**d**; mean ZIP = 8.01), remdesivir and montelukast (**e**; mean ZIP = 13.64), and nirmatrelvir and montelukast (**f** ; mean ZIP = -1.84). Simulations were performed using experimentally determined IC_50_ values and the independently calibrated kinetic model without re-estimation of replication parameters. **g** to **i**: Structural modeling of inhibitor binding to the SARS-CoV-2 main protease (NSP5). **g** Binding mode of nirmatrelvir within the NSP5 catalytic pocket PDB: 8DZ2. **h** Highest-scoring docking pose of montelukast in NSP5, simulated with AutoDock Vina and an adapted version of crystal structure 8DZ2. **i** Superimposed docking poses of nirmatrelvir and montelukast demonstrating substantial overlap within the NSP5 active-site pocket.

## Discussion

By combining high-resolution multi-omics measurements with mechanistic modeling, we established a quantitative framework for describing and predicting intracellular SARS-CoV-2 replication dynamics. The resulting model integrates viral RNA, protein, and infectivity measurements obtained throughout the first 24 h of infection and provides a coherent description of the temporal progression from genome replication to virion release. The model accurately reproduces experimentally observed kinetics and captures key transitions in the viral replication cycle, including RNA amplification, protein accumulation, intracellular virion formation, and extracellular virus release. Beyond reproducing observed data, it enables estimation of internal replication states that are difficult to measure experimentally, including negative-strand RNA intermediates and aggregated NSP dynamics.

The combination of dense time-resolved measurements and mechanistic modeling further enabled the identification of dominant processes governing SARS-CoV-2 replication. Sensitivity analysis revealed that only a small subset of parameters exerts a major influence on viral replication dynamics. In particular, the RNA replication rate (*k_RdRp_*) and the efficiency of NSP maturation (*p_NSP_*) emerged as critical determinants of viral fitness. While *p_NSP_* exerted its strongest influence during the early stages of infection, the impact of *k_RdRp_* persisted throughout the entire simulation period, highlighting viral RNA synthesis as a major bottleneck in productive infection. In contrast, parameters such as ribosome abundance and translation capacity, although highly influential, are unlikely to represent therapeutically tractable targets because they are essential for host-cell viability. Consistent with observations in other positive-strand RNA viruses [19, 20], our analysis also identified DMV-associated processes as important determinants of replication efficiency. In particular, degradation within replication compartments strongly affected viral output, emphasizing the protective role of DMVs in shielding viral RNA from degradation and innate immune recognition [22]. Although no compounds currently applied in the clinics target DMV formation or stability, these findings suggest that replication compartment biology may represent an underexplored avenue for antiviral intervention.

A central objective of this study was to evaluate whether the model could accurately predict viral behavior under perturbations that were not used during model calibration. To this end, we challenged the model with antiviral compounds targeting distinct stages of the replication cycle. Notably, two of the most influential parameters identified by the sensitivity analysis correspond directly to the targets of remdesivir and nirmatrelvir, providing an opportunity for independent validation. Overall, the model reproduced the experimentally observed effects of both compounds with good accuracy, demonstrating that the inferred kinetic structure generalizes beyond the original calibration dataset. Moreover, our results suggest that none of the evaluated drug interventions achieve a sustained reduction in viral load. Instead, treatment primarily alters infection dynamics by delaying the time to peak viral load (S6 Fig.), without changing the overall trajectory of viral replication. This behavior is consistent with clinical observations in which many antiviral therapies reduce viral kinetics or symptom severity but fail to fully prevent viral expansion when administered alone [23]. Our findings therefore support the interpretation that antiviral therapy may be insufficient for effective infection control.

For remdesivir, simulations based on the nominal IC_50_ initially overestimated antiviral efficacy. Improved agreement with experimental observations was obtained when a lower effective intra-cellular inhibition was assumed, suggesting that intracellular drug activation, restricted access to replication complexes within DMVs, and viral proofreading mechanisms may reduce the effective inhibition experienced by the replication machinery [24]. This observation illustrates the importance of considering intracellular drug availability and target engagement when translating biochemical potency into antiviral efficacy.

For nirmatrelvir, experimental inhibition exceeded model predictions. A likely explanation is that inhibition of NSP5 affects multiple downstream processes that are not explicitly represented in the current model. NSP5-dependent processing is required for the maturation of numerous viral proteins, several of which contribute to modulation of host antiviral responses [25]. Consequently, inhibition of NSP5 may produce indirect antiviral effects that extend beyond the reduction of replication activity represented by *p_NSP_*. Since host innate immune responses are not explicitly modeled, such effects cannot currently be reproduced and likely contribute to the observed discrepancy.

Perhaps the strongest demonstration of the model’s utility is its ability to discriminate between competing mechanistic hypotheses. The antiviral mechanism of montelukast remains incompletely understood, with both NSP1- and NSP5-mediated effects having been proposed previously [16–18]. While both mechanisms produced *in-silico* reasonable agreement with single-drug perturbation experiments, only the NSP5-targeting scenario accurately reproduced the *in-vitro* experimentally observed behavior in combination therapy experiments. In particular, the absence of synergy between montelukast and nirmatrelvir was consistent with both compounds acting on the same molecular target. Independent docking simulations further supported this interpretation by indicating overlapping binding sites within the NSP5 catalytic pocket. Together, these findings support NSP5 as the primary antiviral target of montelukast and demonstrate how mechanistic models can be used not only to predict perturbation responses but also to infer underlying biological mechanisms.

Compared with previously published intracellular SARS-CoV-2 replication models [9, 10, 13, 14], the present framework benefits from dense multi-omics measurements spanning multiple stages of the viral replication cycle. The integration of transcriptomic, proteomic, and infectivity data enables resolution of temporal transitions that are difficult to capture using sparsely sampled datasets. More broadly, our results demonstrate the value of mechanistic models as complementary tools to experimental virology. When coupled to sufficiently informative datasets, such models provide a quantitative framework for evaluating antiviral strategies, predicting drug interactions, and generating experimentally testable hypotheses.

Several limitations should be considered. First, the model was calibrated using *in-vitro* data from A549 (A/T) cells infected with the Wuhan strain of SARS-CoV-2. Consequently, parameter values may not directly transfer to other cell types or viral variants and would require reestimation. Second, the model focuses exclusively on infectious virions and does not explicitly represent non-infectious particles, which constitute a substantial fraction of released virus [26]. As a result, some processes, including losses of viral RNA during non-productive replication, are implicitly absorbed into effective parameter estimates. Third, all non-structural proteins are represented as a single aggregated *NSP* state. While this abstraction captures the core functionality of the replication machinery, it does not account for protein-specific activities such as immune evasion or host-cell modulation. Fourth, host immune responses, including interferon signaling and innate antiviral defenses, are not represented explicitly. Finally, the model describes the average behavior of an infected cell and therefore cannot capture cell-to-cell variability that may influence replication kinetics at the population level.

Future extensions should incorporate explicit host responses, increase the resolution of individual viral proteins, and evaluate model transferability across additional cell types and viral variants. Emerging technologies such as single-cell transcriptomics and live-cell RNA tracking may provide the data required to resolve early replication events and stochastic variability in greater detail.

In conclusion, we present a quantitatively calibrated framework for studying intracellular SARS-CoV-2 replication that integrates high-resolution experimental measurements with mechanistic modeling. Beyond reproducing observed replication kinetics, the model identifies critical rate-limiting processes, predicts responses to antiviral perturbations, and enables discrimination between competing mechanistic hypotheses. Given the conserved architecture of CoV replication, the framework should be readily adaptable to additional CoV variants and related viruses, providing a foundation for future studies of viral replication and antiviral intervention strategies.

## Materials and methods

### Measurements of SARS-CoV-2 replication in A549-ACE2 cells

#### Cell culture

A549 cells stably overexpressing human ACE2 and TMPRSS2 (A549 (A/T); [27]) were cultured in Dulbecco’s Modified Eagle Medium (DMEM) supplemented with 5% (v/v) fetal calf serum (FCS), 1% (v/v) non-essential amino acids (NEAA), 100 units/mL penicillin, 100 µg/mL streptomycin, and 2 mM L-glutamine. To maintain selection pressure, cells were continuously cultured in the presence of blasticidin (10 µg/mL) and puromycin (0.5 µg/mL). Vero E6 cells (ATCC CRL-1586) were maintained in DMEM supplemented with 10% (v/v) FCS, 1% (v/v) NEAA, 100 units/mL penicillin, 100 µg/mL streptomycin, and 2 mM L-glutamine. All cell lines were incubated at 37 °C in a humidified atmosphere with 5% CO_2_. Medium was changed every 2–3 days, and cells were passaged at 70–80% confluence using 0.05% trypsin-EDTA. Cells were routinely tested for mycoplasma contamination and confirmed negative.

#### Infection and time-course sampling

For analysis of viral replication kinetics, 5 *×* 10^4^ A549 (A/T) cells were seeded per well in 24-well plates and allowed to adhere at 37 °C in a humidified atmosphere with 5% CO_2_. Cells were infected with SARS-CoV-2 (Wuhan WT, passage 2; GISAID accession ID: EPI_ISL_1118929) at a multiplicity of infection (MOI) of 1 using an inoculum prepared in ice-cold cultivation media. To synchronize infection, cells were incubated with the inoculum at 8 °C for 30 min, followed by an incubation at 37 °C for 1 h. The temperature shift from 8 °C to 37 °C was defined as the starting point of infection (0 hpi). After 1 h, the inoculum was removed, cells were washed three times with PBS and 500 µL of fresh culture medium was added per well. The first samples (1 hpi) were collected immediately thereafter. Samples were collected hourly during the first 8 hpi and every two hours thereafter until 24 hpi. For each time point, supernatants were collected and stored at -80 °C for subsequent titration on VeroE6 cells. Cells were washed three times with PBS prior to downstream processing. For quantification of intracellular viral titers, cells were resuspended in 200 µL PBS and subjected to three freeze/thaw cycles. Cell debris was removed (300 *×* g, 5 min), and the supernatant was used for titration on VeroE6 cells. For RNA isolation, cells were processed using the Qiagen RNeasy Mini Kit according to the manufacturer’s instructions, and RNA samples were stored at -80 °C until further analyses. For protein analysis, cells were lysed in using different buffers depending on the downstream application. For western blot analysis, cells were lysed in M-PER buffer (Thermo Fisher Scientific) according to the manufacturer’s instructions. For LC-MS/MS analysis, proteins were isolated using urea buffer (30 mM Tris HCl, 7 M Urea, 2 M thiourea, 0.1% NaDOC, pH 8.5). Lysates were stored at -80 °C prior to analysis. All experiments were performed with at least three biological replicates.

#### Viral titers

To determine extra- and intracellular virus titers, supernatants were harvested at the respective time points. For intracellular virus quantification, cells were subjected to three freeze/thaw cycles in 200 µL PBS. Viral loads of the samples were then determined by an endpoint dilution assay. Briefly, 1 *×* 10^4^ Vero E6 cells were seeded per well in 96 well plates one day prior to infection. The next day, samples were serially diluted (10-fold serial dilutions) and incubated on the Vero E6 cells for 4 days at 37 °C in a humidified atmosphere with 5% CO. Subsequently, supernatants were removed and the cells were fixed with 4% paraformaldehyde (PFA) in PBS and stained with crystal violet. Cytopathic effects (CPE) were assessed by visual inspection, and median tissue culture infectious dose (TCID_50_) per mL was calculated using the Spearman-Kärber method.

### Transcriptomic analyses

#### RNA isolation and RT-qPCR

For transcriptomic analysis, culture supernatants were removed and cells were washed three times with PBS. Total RNA was isolated using the RNeasy Mini Kit (Qiagen) according to the manufacturer’s instructions, including on-column DNase I digestion (Qiagen). RNA concentrations were measured using a NanoDrop spectrophotometer (Thermo Fisher Scientific). For quantitative analyses, 50 ng of RNA per sample was subjected to a probe-based one-step RT-qPCR using the 1 step RT qPCR Kit (Promega). Primer-probe sets targeting sgRNA E and gRNA were used as previously described [28].

#### Library preparation and sequencing

To validate relative viral RNA expression, and especially sgRNA species of SARS-CoV-2 within the kinetic samples, we used the commercially available, rapid sequencing ARTIC protocol with Oxford Nanopore technology. For kinetic, we preformed library preparation according to protocols developed by the ARTIC Network (Version 3, [29]). In brief, following cDNA synthesis, multiplex PCR was performed using the Varscip short V2 Primer mix I or II. Following purification, we applied 60 ng per sample to the library preparation using the Amplicon barcoding with Native Barcoding Expansion 96 (EXP-NBD196, and SQK-LSK109). Each library pool was loaded onto a Spot-ON Flow Cell (R9.4.1) and sequencing was conducted in MinKNOW for 72 h, resulting in an average read coverage per barcode of 324,257, ranging from 50,404 to 854,197 reads per barcode.

#### Absolute sgRNA count determination

Raw nanopore sequencing reads were processed using Porechop (v0.2.4; [30]) and Cutadapt (v4.1; [31]) for adapter trimming. Read quality was assessed using FASTQC (v0.11.9; [32]). Subsequently, Periscope (v0.1.2; [33]) was employed to determine the sgRNA counts per hundred thousand reads. To obtain absolute sgRNA copy numbers, count values derived from Periscope were normalized to absolute RNA copy numbers determined by RT-qPCR targeting the E-gene. Specifically, the relative abundance of each sgRNA species was scaled based on the corresponding absolute E gene copy number per sample. Finally, the sgRNA counts were normalized to copies per cell.

#### Mass spectrometry

Proteome quantification was carried out as described before [34]. Briefly, cell lysates were processed using the SP3 protocol with slight modifications. 5 µg protein per sample were reduced with dithiothreitol (5 mM, 50 °C, 15 min) and alkylated with 2-iodoacetamide (15 mM, RT, 15 min). After addition of 50 µg SP3-beads and adjustment to 100 µL with 50 mM ammonium bicarbonate (ambic), 170 µL ACN were added and samples were incubated for 18 min. Beads were washed twice with 170 µL 70% EtOH and once with 180 µL ACN, followed by overnight digestion with 0.2 µg trypsin (SERVA Electrophoresis, Heidelberg, Germany) in 36 µL ambic at 37 °C. Samples were dried and reconstituted in 0.1% trifluoroacetic acid. 300 ng tryptic peptides per sample were analyzed in randomized order on a Vanquish Neo UHPLC coupled to an Orbitrap 480 mass spectrometer (both Thermo Scientific). Peptides were trapped on an Acclaim PepMap 100 column (100 µm *×* 2 cm) and separated on a DNV PepMap Neo column (75 µm *×* 150 mm, both Thermo Scientific) using a 1–40% ACN gradient over 120 min at 400 nL/min and 60 °C (mobile phases: 0.1% FA; 80% ACN, 0.1% FA). MS1 spectra were acquired at 350–1450 m/z (resolution 120,000, AGC 300%, IT 54 ms, RF lens 55%). MS2 spectra were recorded at 30,000 resolution with 30% HCD, 145–1450 m/z, AGC 2500%, and IT 80 ms. Forty isolation windows between 350 and 1450 m/z were cycled through with one MS1 scan per 21 MS2 scans. Protein identification and quantification were performed in library-free mode using DIA-NN (v1.8.1) against the SwissProt Homo sapiens database and the UniProt SARS-CoV-2 reference proteome (both v. 2022_01), with the neural network classifier in double-pass mode and species-specific protein inference. Data were filtered for 0.01 FDR and protein matrices were used for further processing.

#### Western blotting

Quantitative western blot analysis was performed to determine the SARS-CoV-2 N expression levels over time. Cells were harvested at the indicated time points from infection experiments as described above. Cells were lysed in 200 µL M-PER buffer (Thermo Fisher Scientific) supplemented with protease inhibitors (Roche). After 5 min incubation, 50 µL Laemmli buffer was added and samples were heat inactivated at 95 °C for 15 min. For each sample, 10 µL of lysate was loaded onto an 8% SDS-PAGE gel. In parallel, a serial dilution of recombinant N (Abcam) was included as a standard curve for absolute quantification. Following gel electrophoresis, samples were transferred onto nitrocellulose membranes (BioRad) using wet transfer. Membranes were blocked for 1 h at room temperature in PBS-T containing 5% milk powder. Blots were incubated overnight at 4 °C with primary antibodies against SARS-CoV-2 N (antibodies-online, 1:2000) and *β*-actin (Fisher Scientific 1:10,000; loading control). After washing three times with PBS-T, membranes were incubated with a horseradish peroxidase-(HRP) conjugated secondary antibody (BIOZOL Diagnostica) for 1 h at room temperature (according to the manufacturer’s instructions). Signal detection was performed using the Pierce ECL Plus Western Blotting Substrate (Thermo Fisher Scientific) and an Azure300 imaging system. Images were analyzed using Fiji software. The Western blot-derived protein abundance data were used for normalization of the mass spectrometry data (see Supporting Text).

### Validation experiments

#### Antiviral dose-response analysis

For validation experiments, the FDA-approved antiviral compounds nirmatrelvir (Hycultec), montelukast (Hycultec) and remdesivir (Carl ROTH) were used. Compounds were dissolved and diluted in DMSO. Serial dilution series were prepared in DMSO prior to experiments and compounds were stored at -80 °C until use. Infection experiments were performed as described above. Compounds were added to the cells 1 h prior to infection. Final DMSO concentrations in the culture medium was 0.1% (v/v) for nirmatrelvir and remdesivir, and 1% for montelukast. Vehicle controls containing corresponding DMSO concentrations were included in all experiments. At 24 hpi, supernatants of the samples were collected and infectious viral titers determined as described above using TCID_50_ assays. Dose-response curves were generated by normalizing viral titers to the respective vehicle controls. IC_50_ values were determined using a four-parameter log-logistic model based on the Hill equation, implemented in GraphPad Prism. All biological replicates were included in the fitting procedure.

#### Time-course analysis at IC_50_ concentrations

Time-course experiments under antiviral treatment were performed as described above, with the modification that compounds were added 1 h prior to infection and maintained throughout the experiment at their respective IC_50_ concentrations. Vehicle control samples were supplemented with the corresponding DMSO concentrations in the culture media.

#### Combination treatment

For combination treatments, respective dilutions series of nirmatrelvir, remdesivir, and montelukast were applied in pairwise combination. DMSO concentrations in the culture medium was 0.1% (v/v) in the case of nirmatrelvir and remdesivir combination, and 1% (v/v) for the other two combinations, which included montelukast. Infection and sampling were performed as described above. Drug interactions were assessed based on dose-response matrices and subsequent analysis using SynergyFinder R package (Version 3.10.3 [35, 36]). The calculation of synergy scores and the generation of corresponding 2D visualizations were executed via a custom R script [37, 38] with minor modifications.

#### Cell viability (MTT assay)

Cell viability was determined by seeding 10^4^ cells per well in a 96 well plate and incubating at 37 °C and 5% CO_2_ overnight. Cells were treated with the respective compound concentrations or DMSO as vehicle control for 24 h. Next, 0.5 mg/mL 3-(4,5-dimethylthiazol-2-yl)-2,5-diphenyltetrazolium bromide (MTT) substrate were applied to the cells and incubated for further 90 min. Medium was then removed and 50 µL of DMSO was added to each well. The absorbance of each well was read on a microplate absorbance reader (Tecan) at 570 nm. Experiment was repeated in three individual replicates. After medium background subtraction, values were normalized to the vehicle control. Cell viability data are shown in Supplementary Fig. S9.

#### Molecular docking simulation

Molecular docking simulations were performed using AutoDock Vina [39, 40] provided by Swiss-Dock [41]. The crystal structure of NSP5 (PDB ID: 8DZ2), co-crystallized with nirmatrelvir, was used as the target. Prior to docking, nirmatrelvir was removed as ligand from the structure and no further modifications were performed. As input for the ligand, we used the SMILES representation of montelukast:

CC(C)(O)C1=CC=CC=C1CC[C@@H](SCC1(CC(O)=O)CC1)C1=CC=CC(\C=C\C2=NC3=C(C=CC(Cl)=C3)C=C2)=C1

Docking was performed within a search space centered on the NSP5 active site (box center: x = -18, y = 5, z = 29; box size: x = 30, y = 30, z = 30). The exhaustiveness parameter was set to 35. In total, 20 fits were provided by AutoDock Vina, whereas the highest scored 10 fits were all in the same grove of the protein. The highest-scoring docking pose was selected for visualization and is shown in Fig. 5.

#### Mathematical model

Our model of the intracellular replication cycle of SARS-CoV-2 is based on the scheme shown in Fig. 1. Import/assembly of virions and translation of non-structural proteins are adapted from a previously published model by Zitzmann et al. [19]. Viral entry is modeled as a non-reversible import process (eqs. (1) and (2)) where *V* represents the extracellular inoculum virus used for infection. *V* is subsequently imported into the cell at a rate of *k_in_* which summarizes binding and entry of the virus and the release of one copy positive-sense gRNA (*R_G_*_(+)_) into the cytoplasm. Given the short time-scale of the experiment and the rapid import, we assume no degradation for *V*, *V_Cyt_* and *V_out_* within the model. This assumption is supported by profile likelihood estimation, which indicates that degradation rates of virions are partially practically identifiable and converge toward negligible values under the current data constraints.

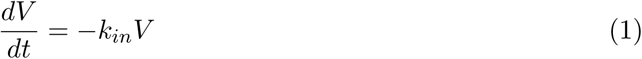

The steps of gRNA translation are described by equations (2–4). Here, *R_G_*_(+)_ denotes positive-strand gRNA present in the cytoplasm of the cell, *TC*_1_*_ab_* the translation complex of the polyprotein ORF1ab, and *NSP* the non-structural proteins that result from the autoproteolysis of the ORF1a and ORF1ab polyproteins. We will only consider the polyprotein ORF1ab to avoid unnecessary complexity in the model. The polyprotein ORF1ab is translated from the gRNA via ribosomal frameshifting whereby ORF1a consists of NSP1–NSP11, while ORF1ab consists of NSP1–NSP16 without NSP11. To reduce model complexity, we do not explicitly resolve individual non-structural proteins. Instead, all NSPs are aggregated into a single effective variable *NSP*, representing the combined functional activity of the viral replication machinery. The corresponding equations for translation are as follows:

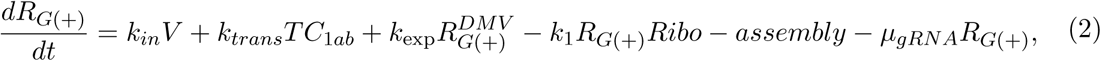

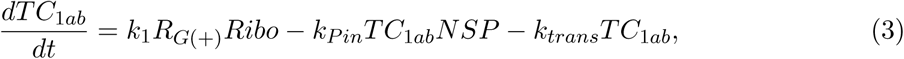

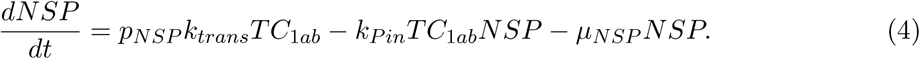

The *assembly* term in equation 2 describes the assembly of gRNA and structural proteins into viral particles. This term is described in more detail in equation 15. Endocytotically imported gRNA *R_G_*_(+)_ binds reversibly with rate *k*_1_ to free ribosomes *Ribo* in the cytoplasm and forms a translation complex *TC*_1_*_ab_* (eq. 3). Assuming the number of total ribosomes *Ribo_tot_*is constant in the cell, the number of free ribosomes for translation can be determined by *Ribo* = *Ribo_tot_ − TC_tot_*, where *TC_tot_* = *TC*_1_*_ab_* + *TC_S_* + *TC_E_* + *TC_N_* + *TC_M_* + *TC*_3_*_a_* + *TC*_6_ + *TC*_7_*_a_* + *TC*_8_ represents the total number of all translation complexes. Then, *TC*_1_*_ab_* translates the non-structural proteins *NSP* at a rate *k_trans_*, which are subsequently imported with *TC*_1_*_ab_* into the DMV at a rate *k_P in_* and form a replication complex *RC* (eq. 5). To simulate the inhibition of NSP5, we included *p_NSP_*which reflects the probability to generate functional NSP proteins via NSP5 induced autoproteolysis. The ribosome of *TC*_1_*_ab_* stays in the cytoplasm. The degradation of *NSP* and *R_G_*_(+)_ takes place with degradation constants *µ_NSP_* and *µ_gRNA_*. However, not all *R_G_*_(+)_ are used for translation. Some are used for virus assembly, which is described in eq. 16 below.

The discontinuous transcription mechanism of CoVs is unique among viruses and is described below with the eqs. (5-10). Due to this mechanism, a large variety of different RNAs are produced during replication. Let *i ∈ {G, S, E, M, N,* 3*a,* 6, 7*a,* 8*}* denote the different RNA species for gRNA (*G*), the structural proteins spike *S*, envelope *E*, membrane *M*, nucleocapsid *N*, and accessory proteins ORF3a 3*a*, ORF6 6, ORF7a 7*a*, and ORF8 8. Notably, the replisome, consisting of multiple NSPs, is simplified into *NSP^DMV^* to reduce model complexity. *RC* describes the replication complex for the production of a nested set of negative (s)gRNAs *R_i_*_(_*_−_*_)_, which are transcribed at *RC_i_* into a set of positive sgRNAs 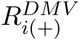 via *NSP^DMV^*. The sgRNAs are subsequently exported into the cytoplasm, while the gRNA can undergo another replication cycle.

*RC* produces negative RNAs *R_i_*_(_*_−_*_)_ at a rate *k*_2_*_i_* = *k_RdRp_/l_i_* and probability *p_i_*. The negative RNAs form additional replication complexes *RC_i_*_(+)_ with on *NSP^DMV^* at rate *k*_3_ (eq. 7), which in turn produce positive (s)gRNAs 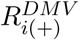 at rate *k*_2_*_i_* (eqs. 8 and 9).

The positive sgRNAs of the proteins *P ∈ {S, E, M, N,* 3*a,* 6, 7*a,* 8*}* are exported from the replication vesicle at a rate *k_exp_* (eq. 9). The positive gRNA 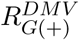 can either be exported into the cytoplasm at a rate *k_exp_* or form a new replication complex *RC* with *NSP^DMV^* at a rate *k*_4_ (eqs. 8 and 10). All species in the DMV are degraded with the same degradation rate *µ_DMV_*.

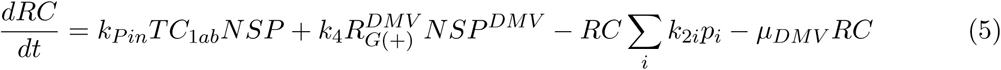

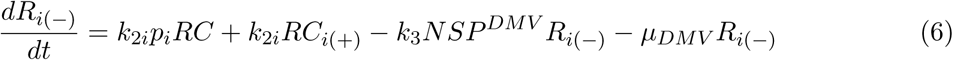

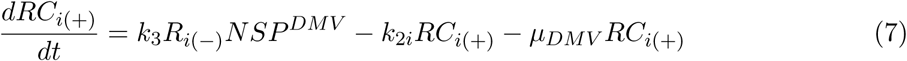

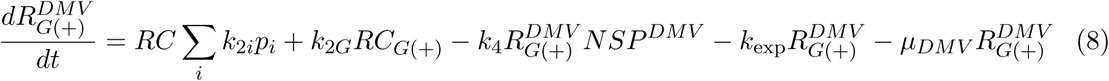

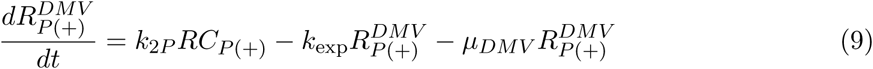

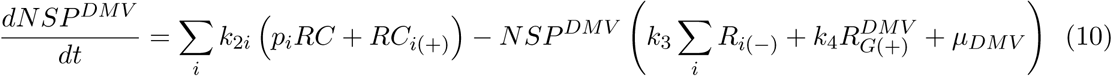

The exported positive RNAs *R_P_* _(+)_ of the proteins *P* form translation complexes *TC_P_* with free ribosomes *Ribo*, which then translates the respective structural and accessory proteins (eqs. 11-14). Here, similarly to the translation of ORF1ab, positive sgRNAs 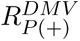 binding to the free ribosomes *Ribo* at a rate *k*_1_ (eq. 12). These translate the respective structural proteins *SP ∈ {S, E, M, N }* and accessory proteins *AP ∈ {*3*a,* 6, 7*a,* 8*}* at a rate *k_trans_*(eqs. 13 and 14). The structural proteins are then used for virus assembly, while the accessory proteins are not modeled as feeding back on viral replication dynamics for simplicity of the model. All proteins and sgRNAs degrade at a degradation rate of *µ_P_* and *µ_RNA_*.

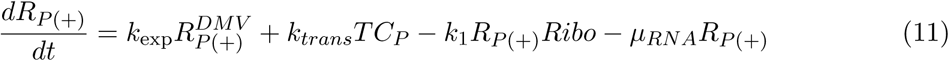

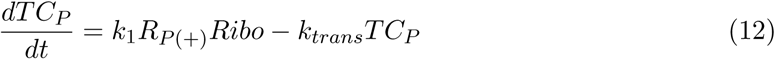

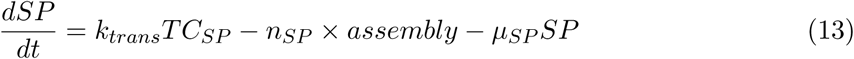

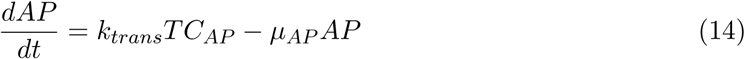

The assembly and export of the viruses proceed analogously to those described in the Zitzmann et al. model [19] (eqs. 15-17).

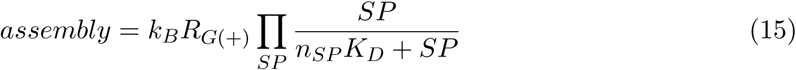

*n_SP_*represents the number of structural proteins *SP* required for a virion and *K_D_*is an Michaelis–Menten-like half-saturation constant for structural protein-dependent virion assembly. If sufficient structural proteins *SP* and a copy of the gRNA *R_G_*_(+)_ are present in the cytoplasm, they are assembled with rate *k_B_* into new infectious virus particles *V_Cyt_* (eq. 16). The assembled viruses are subsequently exported from the cell at a rate *k_out_* (eqs. 16 and 17).

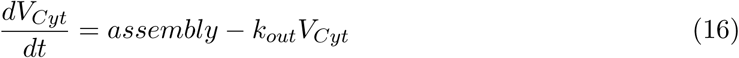

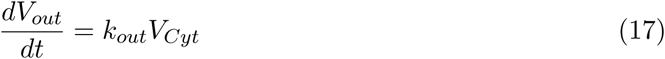

#### Model assumptions

To simplify the complex intracellular replication cycle of SARS-CoV-2, we made the following assumptions. The model itself represents an average cell. Consequently, some molecular species may have fractional molecule numbers (i.e. <1 copy) over the course of the simulation. We modeled only a single double-membrane vesicle (DMV) based on Dahari et al. [42]. All species associated with the DMV degrade at the same rate. To reduce the number of equations and keep the model simple, we aggregated all NSP proteins into a single *NSP* variable. We do not explicitly distinguish between ORF1a and ORF1ab translation and instead represent NSP production through a single ORF1ab-derived pathway. Double-stranded replication intermediates are not explicitly represented. We included only canonical ORF proteins and RNAs in the model, non-canonical ORFs and ORF10 were not included because of their low abundance and the lack of quantitative measurements. The model itself tracks only infectious virions, as defective particles were not quantified experimentally. Virion degradation was not included over the 24 h observation period, as supported by parameter identifiability analysis. A single degradation rate for all sgRNAs is applied, to reduce the total number of parameters and model complexity. We also do not incorporate direct cellular immune response in the equations. Drug treatment simulations assume that intracellular drug concentrations remain sufficiently stable over the course of the experiment to maintain a constant inhibitory effect.

#### Parameter estimation

The model was implemented in MATLAB version 24.2.0 (R2024b) [43] and consists of 49 parameters, of which 28 were fixed based on literature evidence, calculations, and optimization processes (see Table 1) and the remaining parameters were estimated from data. For the optimization of the log-likelihood function we used a trust-region algorithm combined with a Latin hypercube approach within the toolbox Data2Dynamics [44, 45]. In total, 1000 independent multi-start fitting procedures were performed. For each fitting procedure, 100 initial parameter vectors were generated by Latin hypercube sampling from the predefined parameter bounds, resulting in 100,000 local optimization runs in total. This strategy was used to explore a wide range of parameter combinations, thereby increasing the likelihood of convergence to the global minimum. Experimental data were converted to comparable cellular quantities before fitting. RNA measurements were expressed as copies per cell, and protein measurements were expressed as molecules per cell. Infectious virus titers were linked to the model-predicted number of virions per cell using an additional observation scaling factor *f Scale_V irus_*. All experimental measurements were log_10_-transformed before fitting. Model calibration was based on the mean of the replicate measurements at each time point rather than on individual replicate values. To calibrate the model, we minimized the negative log likelihood [45]: *L̂* = 2 log(*L*(*ŷ|θ*)), with

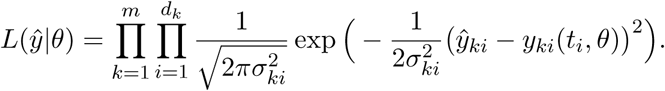

**Table 1:** Overview of all parameters used in the ODE model.

| Parameter | Definition | Value | Unit | 95% CI<br>(log<sub>10</sub>) | Reference |
| --- | --- | --- | --- | --- | --- |
| $l_G$ | Length of ± gRNA | 29,903 | nt | - | NCBI ref seq:<br>NC_045512.2<br>and SI |
| $l_S$ | Length of ± S RNA | 8384 | nt | - | NCBI ref seq:<br>NC_045512.2<br>and SI |
| $l_{3a}$ | Length of ± 3a RNA | 4554 | nt | - | NCBI ref seq:<br>NC_045512.2<br>and SI |
| $l_E$ | Length of ± E RNA | 3702 | nt | - | NCBI ref seq:<br>NC_045512.2<br>and SI |
| $l_M$ | Length of ± M RNA | 3424 | nt | - | NCBI ref seq:<br>NC_045512.2<br>and SI |
| $l_6$ | Length of ± ORF6<br>RNA | 2745 | nt | - | NCBI ref seq:<br>NC_045512.2<br>and SI |
| $l_{7a}$ | Length of ± ORF7a<br>RNA | 2553 | nt | - | NCBI ref seq:<br>NC_045512.2<br>and SI |
| $l_8$ | Length of ± ORF8<br>RNA | 2053 | nt | - | NCBI ref seq:<br>NC_045512.2<br>and SI |
| $l_N$ | Length of ± N RNA | 1673 | nt | - | NCBI ref seq:<br>NC_045512.2<br>and SI |
| $k_{in}$ | Import rate (virions<br>into cell) | 0.431 | h<sup>-1</sup> | - | See methods |
| $k_1$ | $TC$ assembly rate | 0.011112 | Cell molecules<sup>-1</sup><br>h<sup>-1</sup> | [-2.287, -1.503] | |
| $k_{RdRp}$ | Transcription rate | 936,000 | nt h<sup>-1</sup> | - | 260 nt/s [49] |
| $k_{Pin}$ | Import rate (into<br>DMVs) | 9995.9 | Cell molecules<sup>-1</sup><br>h<sup>-1</sup> | [2.13, Inf] | |
| $k_3$ | $RC_i$ assembly rate | 1000 | Cell molecules<sup>-1</sup><br>h<sup>-1</sup> | [1.883, Inf] | |
| $k_4$ | Assembly rate $RC$ | 0.001 | Cell molecules<sup>-1</sup><br>h<sup>-1</sup> | [-Inf, 0.9779] | |
| $p_G$ | Transc. prob. gRNA | 0.0651517 | - | - | See methods |
| $p_S$ | Transc. prob. S RNA | 0.61333 | - | [-0.2944,<br>-0.1684] | |
| $p_M$ | Transc. prob. M RNA | 0.14553 | - | [-0.9735,<br>-0.7121] | |
| $p_N$ | Transc. prob. N RNA | 0.094508 | - | [-1.062,<br>-0.9324] | |
| $p_E$ | Transc. prob. E RNA | 0.0033834 | - | [-2.639, -2.306] | |
| $p_{3a}$ | Transc. prob. ORF3a RNA | 0.02759 | - | [-1.682, -1.443] | |
| $p_6$ | Transc. prob. ORF6 RNA | 0.049292 | - | [-1.5, -1.12] | |
| $p_{7a}$ | Transc. prob. ORF7a RNA | 0.00095522 | - | [-3.135, -2.909] | |
| $p_8$ | Transc. prob. ORF8 RNA | 0.00025968 | - | [-3.716, -3.419] | |
| $k_{exp}$ | DMV export rate (sgRNA) | 280.31 | $h^{-1}$ | [2.236, 3.032] | |
| $k_{trans}$ | Translation rate | 360 | $h^{-1}$ | - | See methods |
| $p_{NSP}$ | Prob. to generate functional NSP | 1 | - | - | See methods |
| $K_D$ | Structure protein assembly saturation constant | 1799.7 | Molecules cell$^{-1}$ | [-0.466, 3.459] | |
| $k_B$ | Assembly rate virion | 9861.9 | $h^{-1}$ | [-1.289, Inf] | |
| $k_{out}$ | Export rate virion | 0.044909 | $h^{-1}$ | [-1.48, -1.219] | |
| $n_E$ | E protein per virion | 10 | - | - | [57] |
| $n_M$ | M protein per virion | 2006 | - | - | [57] |
| $n_N$ | N protein per virion | 456 | - | - | [53, 54] |
| $n_S$ | S protein per virion | 78 | - | - | [50–52] |
| $Virus_{init}$ | Initial number of virions | 1 | Virion cell$^{-1}$ | - | See methods |
| $Ribo_{init}$ | Number of ribosomes | 11,058 | Molecules cell$^{-1}$ | [3.961, 4.139] | |
| $fScale_{Virus}$ | Scaling factor (virus) | 3.7706 | Molecules cell$^{-1}$ TCID$_{50}^{-1}$ | [0.4145, 0.739] | |
| $\mu_{DMV}$ | Degradation rate (inside DMV) | 0.85202 | $h^{-1}$ | [-0.4211, 0.3128] | |
| $\mu_{RNA}$ | Degradation rate (sgRNA) | 590.29 | $h^{-1}$ | [2.279, 3.323] | |
| $\mu_{gRNA}$ | Degradation rate (gRNA) | 3.8302 | $h^{-1}$ | [0.1843, 1.211] | |
| $\mu_{3a}$ | Degradation rate (3a protein) | 0.2596 | $h^{-1}$ | - | [60] |
| $\mu_6$ | Degradation rate (6 protein) | 0.7374 | $h^{-1}$ | - | [60] |
| $\mu_{7a}$ | Degradation rate (7a protein) | 0.023 | $h^{-1}$ | - | 30h estimated [65] |
| $\mu_8$ | Degradation rate (8 protein) | 1.44406 | $h^{-1}$ | - | [60] |
| $\mu_N$ | Degradation rate (N protein) | 0.023 | $h^{-1}$ | - | 30h estimated [66] |
| $\mu_M$ | Degradation rate (M protein) | 0.0866 | $h^{-1}$ | - | 8h estimated [60] |
| $\mu_S$ | Degradation rate (S protein) | 0.023 | $h^{-1}$ | - | 30h estimated, see methods |
| $\mu_E$ | Degradation rate (E protein) | 0.1768 | $h^{-1}$ | - | [60] |
| $\mu_{NSP}$ | Degradation rate (NSP proteins) | 0.12258 | $h^{-1}$ | - | [60] |

The objective is to minimize the distance between the observation *y* and experimental data *ŷ* at time points *t_i_* with *i* = 1*, … , d_k_*. *d_k_* represents the number of experimental data points, with *k* = 1*, … , m*, where *m* denotes the number of observations. 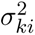 represents the variance for each experimental data point, which were estimated from the biological replicates, therefore residuals were normalized by the experimentally observed variability, giving less weight to data points with higher biological variability and preventing observables with different scales or variances from disproportionately dominating the fit.

Estimated parameters can be found in Table 1. The first 3–4 h of transcriptomic and proteomic measurements were excluded from parameter estimation because the observed signals were dominated by declining input-derived material rather than de novo viral synthesis. Based on literature we fixed as many parameters as possible to simplify the fitting procedure and ensure identifiability for the estimated parameters.

Given an MOI of 1, the expected number of infecting virions per cell is one. For the deterministic ODE model, we therefore initialized the system with *V_init_* = 1. We also initialized the number of total ribosomes based on the estimated parameter *Ribo_init_*. All other initial state conditions are initialized to zero. We fixed the translation rate parameter *k_trans_* under the assumption that ribosomes occupy approximately one binding site per 80 AA [46] on sgRNA and with an average translation elongation rate of 3–8 AA s^-1^ in eukaryotes [47, 48], thus *k_trans_* = 360 h^-1^ with an assumed translation rate of 8 AA s^-1^ = 28,800 AA h^-1^.

Under untreated conditions, *p_NSP_*was fixed to 1, corresponding to fully efficient generation of functional NSPs. Drug treatment simulations modeled NSP5 inhibition by reducing this parameter. Transcription rates *k*_2_*_i_* for RNA species *i* are defined as *k*_2_*_i_* = *k_RdRp_/l_i_* where *l_i_* donates the length of RNA *i*. This assumes a constant elongation rate of the polymerase with an average rate of *k_RdRp_* = 936, 000 nt h^-1^, which is based on a previous study [49] that reported a complete genome replication within 2 min. All RNA lengths *l_i_* are calculated in SI (see S3 Table) and we assume same lengths for corresponding positive and negative strand RNAs.

We introduced parameters *p_i_*, representing the probability of synthesizing RNA species *i* during the discontinuous transcription of SARS-CoV-2 with Σ*_i_p* = 1. To satisfy the normalization constraint, *p_G_* to generate an gRNA was defined as 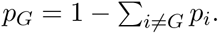

The number of structural proteins *n_SP_* for virus assembly is based on estimates from literature. We assume an average of 26 spike protein trimers per virus. This follows from the studies [[50]; S_3_ = 26 *±* 15], [[51]; S_3_ = 24 *±* 9] and [[52]; S_3_ = 48 with a range of 25–127]. As each trimer consists of three monomers, the total number of spike proteins per virion is fixed as *n_s_* = 3 *×* 26 = 78. The nucleocapsid protein packages the gRNA into around 38 ribonucleoprotein particles (RNP) [53], consisting each of 12 N proteins [54]. Therefore *n_N_* = 38 *×* 12 = 456. The membrane protein exists as a dimer and is the most abundant structural protein in a virion [55]. Published estimates for SARS-CoV, Murine hepatitis virus (MHV), and Feline CoV are around 1100 M dimers [56] and simulations from [57] indicate a stable virus using 1003 M dimers for SARS-COV-2, thus we fix *n_M_*= 1003 *×* 2 = 2006 monomers per virion. The envelope protein forms pentamers that are present in only a few copies per virion [56, 58]. DeDiego et al. report about 20 copies for MHV [59]. Following Pezeshkian et al. [57], we assumed 2 pentameric pores per virion yielding *n_E_*= 2 *×* 5 = 10 monomers.

The degradation rates of the individual structural and accessory proteins were fixed based on literature values. Membrane, nucleocapsid and ORF3a proteins exhibit long half-lives [60] leading to low estimates for their degradation rates. Given its long half-life, the spike protein was assumed to have a half-life of approximately 30 h, comparable to ORF7a and N [60]. As the model represents all NSPs with a single variable, we averaged reported half-lives [60] to obtain a mean half-life of *∼* 5.56 h, leading to a degradation rate of *µ_NSP_≈* 0.12 h^-1^. However, as NSP11 is a small protein of 13 AA, which cannot be detected by regular mass spectrometry, we disregarded it for the calculation.

Based on binding assay kinetics and literature values [61–64], viral entry was constrained to a rate constant *k_in_*within [0.06, 0.43] h^-1^. Because *k_in_*consistently converged toward the upper bound during optimization and was not practically identifiable, it was fixed at 0.43 h^-1^ for all subsequent analyses. The remaining 21 model parameters were estimated by fitting to the experimental data.

Parameter identifiability was assessed by profile likelihood estimation. Parameters with confidence intervals bounded on both sides were classified as identifiable, whereas parameters with an unbounded confidence interval in at least one direction were classified as partially identifiable (Table 1). Accordingly, infinite confidence interval bounds indicate partial identifiability within the current model structure (*k*_3_*, k*_4_*, k_P in_, k_B_*).

Although *µ_RNA_*was identifiable by profile likelihood analysis, its fitted value corresponds to an apparent RNA half-life of approximately 4.5 s and is therefore unlikely to represent the true biochemical degradation rate of SARS-CoV-2 sgRNA. This high effective loss rate likely compensates for processes not explicitly represented in the current model structure, including host-mediated antiviral responses, changes in RNA detectability, or compartmental redistribution. In addition, the model assumes rapid RNA export from double-membrane vesicles, with the same export process also contributing to genomic RNA export. This structural coupling may lead to an overrepresentation of cytosolic or detectable sgRNA, which is compensated by a high effective sgRNA loss rate. Thus, *µ_RNA_* is identifiable within the current model but should not be interpreted as a biologically realistic sgRNA degradation rate.

#### Sensitivity and identifiability analysis

A global sensitivity analysis was performed using eFAST implemented in MATLAB [67]. This method quantifies the influence of individual model parameters on model outputs by calculating first-order and total-order sensitivity indices. Statistical significance was assessed by comparing estimated sensitivity indices against those obtained for a dummy parameter that does not participate in the model equations.

Sensitivity indices were calculated for the following model outputs: intracellular virus *V_Cyt_*, extracellular virus *V_out_*, and total positive-sense gRNA defined as *R_tot_* = *R_G_*_(+)_ + *TC*_1_*_ab_* + 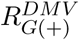 + *RC*. Analyses were performed at an early (3 hpi) and a late (24 hpi) time point to capture temporal differences in parameter influence. Total order sensitivity indices for these parameters are shown in Fig. 3.

Additionally, a profile likelihood analysis (PLA) was performed to asses practical parameter identifiability of all fitted model parameters and to identify parameter dependencies within the fitted model (S10 Fig.). This approach systematically varies one parameter while re-optimizing all remaining parameters to minimize the objective function, thereby assessing whether changes in one parameter can be compensated by adjustments in others. Ninety-five percent confidence intervals were derived from the resulting profile likelihood curves (Table 1). Finite confidence intervals were interpreted as evidence of practical parameter identifiability.

#### Simulation of antiviral intervention

Antiviral effects were implemented using a pharmacological Hill function to modulate the effective rates of targeted viral processes. The residual activation function is defined as:

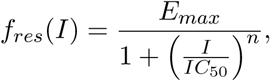

where *I* denotes the drug concentration, *IC*_50_ the half-maximal inhibitory concentration, the Hill coefficient *n* (here *n* = 1) and *E_max_* = 1 the maximal effect of the drug. We fixed *E_max_* = 1 so that the target process retained full activity in the absence of a drug. Drug perturbation experiments were performed using remdesivir, nirmatrelvir, and montelukast. Because cells were pre-treated prior to infection, the residual activation function is applied throughout the full-time course of the experiment, starting at 0 h.

Remdesivir is a nucleotide analogue that causes premature termination of nascent viral RNA synthesis. In the model, this effect was represented by reducing the effective RNA polymerase elongation rate:

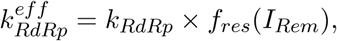

leading to a reduced production of functional RNAs.

The second drug, nirmatrelvir, binds to the active site of NSP5, thereby inhibiting NSP5-mediated proteolytic processing of ORF1a and ORF1ab. The parameter *p_NSP_* represents the probability of generating functional NSPs and was therefore used to model NSP5 inhibition. When multiplying *p_NSP_*by *f* (*I_Nir_*), the rate at which effective NSP is produced is now reduced.

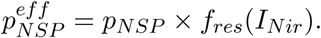

The mechanism of action of montelukast is ambiguous, therefore we tested two different described drug targets independently of each other: NSP5, analogous to nirmatrelvir, and NSP1. NSP1 modulates host ribosome function and promotes preferential translation of viral transcripts. Within the model, this process is represented by the translation initiation parameter *k*_1_. Thus

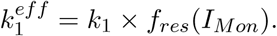

The simulations were performed in R [68]. For the combined drug treatments, as shown in Fig. 5, we inhibited both parameters simultaneously by multiplying them with the corresponding inhibition functions. To simulate synergistic interactions on a same target, we applied the activation function twice:

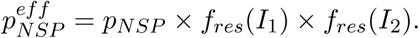

To simulate non-synergistic interactions between two compounds at the same target, we only considered the most effective drug:

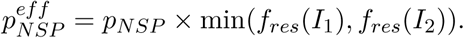

Synergy scores for the combination treatments were computed in R using the SynergyFinder package [35–38], based on the simulated drug inhibition and experimental data.

## Acknowledgements

We acknowledge the valuable input and constructive discussions provided by members of the Department for Molecular and Medical Virology of the Ruhr University Bochum, the Research Unit Emerging Viruses (LIV) and the Institute of Bioinformatics (UMG). We also thank the Department for Molecular and Medical Virology of the Ruhr University Bochum and Institute of Virology, University Hospital Essen, University Duisburg-Essen, Essen, Germany for the usage of high-containment facilities.

## Author contributions

S.T.H., S.W., N.H., T.L.M. and T.B. conducted the experiments, and T.K. performed the mathematical modeling and predictions. T.K., S.T.H., and L.D.B. prepared the first draft of the manuscript. T.K. and S.T.H. prepared the figures. B.S. provided resources. S.P. and L.K. conceived the idea for the study and supervised the project. All authors contributed to revising the manuscript and approved the final version for publication.

## Data availability statement

All processed data supporting the findings of this study, including the numerical data underlying the figures and the parameter estimation results, will be deposited in a public repository and made publicly available upon publication. Data and code will be made available to editors and reviewers upon request during peer review.

## References

[1] V’Kovski, P., Kratzel, A., Steiner, S., Stalder, H. & Thiel, V. Coronavirus biology and replication: implications for SARS-CoV-2. Nat. Rev. Microbiol. 19, 155–170 (2021).

[2] Brant, A. C., Tian, W., Majerciak, V., Yang, W. & Zheng, Z. M. SARS-CoV-2: from its discovery to genome structure, transcription, and replication. Cell Biosci. 11, 136 (2021).

[3] Ryu, G. & Shin, H. W. SARS-CoV-2 infection of airway epithelial cells. Immune Netw. 21, e3 (2021).

[4] Westhoven, S. et al. From zoonotic spillover to endemicity: the broad determinants of human coronavirus tropism. mBio, e0243725 (2025).

[5] Kapischke, T., Herrmann, S. T., Bertzbach, L. D., Pfaender, S. & Kaderali, L. Ordinary differential equation models of SARS-CoV-2 replication dynamics and antiviral drug efficacies. npj Viruses 4 (2026).

[6] Steiner, S. et al. SARS-CoV-2 biology and host interactions. Nat. Rev. Microbiol. 22, 206– 225 (2024).

[7] Qu, B. et al. TMPRSS2-mediated SARS-CoV-2 uptake boosts innate immune activation, enhances cytopathology, and drives convergent virus evolution. Proc. Natl Acad. Sci. USA 121, e2407437121 (2024).

[8] Sergio, M. C., Ricciardi, S., Guarino, A. M., Giaquinto, L. & De Matteis, M. A. Membrane remodeling and trafficking piloted by SARS-CoV-2. Trends Cell Biol. 34, 785–800 (2024).

[9] Ayadi, A. et al. Combining DEVS and semantic technologies for modeling the SARS-CoV-2 replication machinery. IEEE, 1–12 (2021). 10.23919/annsim52504.2021.9552040

[10] Castle, B. T. et al. Biophysical modeling of the SARS-CoV-2 viral cycle reveals ideal antiviral targets. bioRxiv (2020). 10.1101/2020.05.22.111237

[11] Fatehi, F., Bingham, R. J., Dykeman, E. C., Stockley, P. G. & Twarock, R. Comparing antiviral strategies against COVID-19 via multiscale within-host modelling. R. Soc. Open Sci. 8, 210082 (2021).

[12] Isea, R. & Mayo-García, R. First analytical solution of intracellular life cycle of SARS-CoV-2. J. Model Based Res. 1, 6–12 (2022).

[13] Sazonov, I., Grebennikov, D., Meyerhans, A. & Bocharov, G. Sensitivity of SARS-CoV-2 life cycle to IFN effects and ACE2 binding unveiled with a stochastic model. Viruses 14 (2022).

[14] Grebennikov, D. et al. Intracellular life cycle kinetics of SARS-CoV-2 predicted using mathematical modelling. Viruses 13 (2021).

[15] Xu, Z. et al. More or less deadly? A mathematical model that predicts SARS-CoV-2 evolutionary direction. Comput. Biol. Med. 153, 106510 (2023).

[16] Ahmed, M. S. et al. FDA approved drugs with antiviral activity against SARS-CoV-2: from structure-based repurposing to host-specific mechanisms. Biomed. Pharmacother. 162, 114614 (2023).

[17] Durdagi, S. et al. The neutralization effect of montelukast on SARS-CoV-2 is shown by multiscale in silico simulations and combined in vitro studies. Mol. Ther. 30, 963–974 (2022).

[18] Afsar, M. et al. Drug targeting NSP1-ribosomal complex shows antiviral activity against SARS-CoV-2. eLife 11 (2022).

[19] Zitzmann, C. et al. A coupled mathematical model of the intracellular replication of dengue virus and the host cell immune response to infection. Front. Microbiol. 11, 725 (2020).

[20] Binder, M. et al. Replication vesicles are load- and choke-points in the Hepatitis C virus lifecycle. PLoS Pathog. 9, e1003561 (2013).

[21] Noske, G. D. et al. Structural basis of nirmatrelvir and ensitrelvir activity against naturally occurring polymorphisms of the SARS-CoV-2 main protease. J. Biol. Chem. 299 (2023).

[22] Kasuga, Y., Zhu, B., Jang, K. J. & Yoo, J. S. Innate immune sensing of coronavirus and viral evasion strategies. Exp. Mol. Med. 53, 723–736 (2021).

[23] Gonçalves, A. et al. Timing of antiviral treatment initiation is critical to reduce SARS-CoV-2 viral load. CPT Pharmacometrics Syst. Pharmacol. 9, 509–514 (2020).

[24] Agostini, M. L. et al. Coronavirus susceptibility to the antiviral remdesivir (GS-5734) is mediated by the viral polymerase and the proofreading exoribonuclease. mBio 9 (2018).

[25] Gao, Y. & Zhang, J. The role of SARS-CoV-2 main protease in innate immune regulation: from molecular mechanisms to therapeutic implications. Acta Pharm. Sin. B 15, 4497–4510 (2025).

[26] Sender, R. et al. The total number and mass of SARS-CoV-2 virions. Proc. Natl Acad. Sci. USA 118 (2021).

[27] Synowiec, A. et al. Identification of cellular factors required for SARS-CoV-2 replication. Cells 10 (2021).

[28] Roesmann, F. et al. Comparison of the Ct-values for genomic and subgenomic SARS-CoV-2 RNA reveals limited predictive value for the presence of replication competent virus. J. Clin. Virol. 165, 105499 (2023).

[29] Quick, J. nCoV-2019 sequencing protocol v3 (LoCost) v.3. (2020). 10.17504/protocols.io.bp2l6n26rgqe/v3

[30] Wick, R. R., Judd, L. M., Gorrie, C. L. & Holt, K. E. Completing bacterial genome assemblies with multiplex MinION sequencing. Microb. Genom. 3, e000132 (2017).

[31] Martin, M. Cutadapt removes adapter sequences from high-throughput sequencing reads. EMBnet J. 17 (2011).

[32] Andrews, S. FastQC: A quality control tool for high throughput sequence data. (2010). https://www.bioinformatics.babraham.ac.uk/projects/fastqc/

[33] Parker, M. D. et al. Subgenomic RNA identification in SARS-CoV-2 genomic sequencing data. Genome Res. 31, 645–658 (2021).

[34] Maassen, F. et al. The human cytomegalovirus-encoded pUS28 antagonizes CD4+ t cell recognition by targeting CIITA. eLife 14 (2025).

[35] Yadav, B., Wennerberg, K., Aittokallio, T. & Tang, J. Searching for drug synergy in complex dose-response landscapes using an interaction potency model. Comput. Struct. Biotechnol. J. 13, 504–513 (2015).

[36] Zheng, S. et al. Synergyfinder plus: toward better interpretation and annotation of drug combination screening datasets. Genomics Proteomics Bioinformatics 20, 587–596 (2022).

[37] Gömer, A. et al. Dynamic evolution of the sofosbuvir-associated variant A1343V in hevinfected patients under concomitant sofosbuvir-ribavirin treatment. JHEP Rep. 6, 100989 (2024).

[38] Klöhn, M. et al. The glutamate receptor antagonist ifenprodil inhibits Hepatitis E virus infection. Antimicrob. Agents Chemother. 68, e0103524 (2024).

[39] Eberhardt, J., Santos-Martins, D., Tillack, A. F. & Forli, S. AutoDock Vina 1.2.0: new docking methods, expanded force field, and python bindings. J. Chem. Inf. Model. 61, 3891–3898 (2021).

[40] Trott, O. & Olson, A. J. AutoDock Vina: improving the speed and accuracy of docking with a new scoring function, efficient optimization, and multithreading. J. Comput. Chem. 31, 455–461 (2010).

[41] Bugnon, M., et al. SwissDock 2024: major enhancements for small-molecule docking with attracting cavities and AutoDock Vina. Nucleic Acids Res. 52, W324–W332 (2024).

[42] Dahari, H., Ribeiro, R. M., Rice, C. M. & Perelson, A. S. Mathematical modeling of subgenomic Hepatitis C virus replication in Huh-7 cells. J. Virol. 81, 750–760 (2007).

[43] The MathWorks Inc. MATLAB version: 9.13.0 (R2022b). The MathWorks Inc. (2022). https://www.mathworks.com

[44] Raue, A. et al. Data2Dynamics: a modeling environment tailored to parameter estimation in dynamical systems. Bioinformatics 31, 3558–3560 (2015).

[45] Raue, A. et al. Lessons learned from quantitative dynamical modeling in systems biology. PLoS One 8, e74335 (2013).

[46] Kim, H. & Yin, J. Robust growth of human immunodeficiency virus type 1 (HIV-1). Biophys. J. 89, 2210–2221 (2005).

[47] Lodish, H. F. & Jacobsen, M. Regulation of hemoglobin synthesis. J. Biol. Chem. 247, 3622–3629 (1972).

[48] Palmiter, R. D. Ovalbumin messenger ribonucleic acid translation. J. Biol. Chem. 248, 2095–2106 (1973).

[49] Campagnola, G., Govindarajan, V., Pelletier, A., Canard, B. & Peersen, O. B. The SARS-CoV NSP12 polymerase active site is tuned for large-genome replication. J. Virol. 96, e0067122 (2022).

[50] Yao, H. et al. Molecular architecture of the SARS-CoV-2 virus. Cell 183, 730–738 e13 (2020).

[51] Ke, Z. et al. Structures and distributions of SARS-CoV-2 spike proteins on intact virions. Nature 588, 498–502 (2020).

[52] Klein, S. et al. SARS-CoV-2 structure and replication characterized by in situ cryo-electron tomography. Nat. Commun. 11, 5885 (2020).

[53] Zhao, H. et al. Assembly of SARS-CoV-2 nucleocapsid protein with nucleic acid. Nucleic Acids Res. 52, 6647–6661 (2024).

[54] Carlson, C. R. et al. Reconstitution of the SARS-CoV-2 ribonucleosome provides insights into genomic RNA packaging and regulation by phosphorylation. J. Biol. Chem. 298, 102560 (2022).

[55] Zhang, Z. et al. Structure of SARS-CoV-2 membrane protein essential for virus assembly. Nat. Commun. 13, 4399 (2022).

[56] Neuman, B. W. et al. A structural analysis of M protein in coronavirus assembly and morphology. J. Struct. Biol. 174, 11–22 (2011).

[57] Pezeshkian, W. et al. Molecular architecture and dynamics of SARS-CoV-2 envelope by integrative modeling. Structure 31, 492–503 e7 (2023).

[58] Rahman, M. S. et al. Mutational insights into the envelope protein of SARS-CoV-2. Gene Rep. 22, 100997 (2021).

[59] DeDiego, M. L. et al. A severe acute respiratory syndrome coronavirus that lacks the E gene is attenuated in vitro and in vivo. J. Virol. 81, 1701–1713 (2007).

[60] Li, W., et al. Stability of SARS-CoV-2-encoded proteins and their antibody levels correlate with interleukin 6 in COVID-19 patients. mSystems 7, e0005822 (2022).

[61] Baggen, J. et al. Genome-wide CRISPR screening identifies TMEM106B as a proviral host factor for SARS-CoV-2. Nat. Genet. 53, 435–444 (2021).

[62] Ozono, S. et al. SARS-CoV-2 D614G spike mutation increases entry efficiency with enhanced ACE2-binding affinity. Nat. Commun. 12, 848 (2021).

[63] Walls, A. C. et al. Structure, function, and antigenicity of the SARS-CoV-2 spike glycoprotein. Cell 181, 281–292 e6 (2020).

[64] Zhu, Y., Yu, D., Yan, H., Chong, H. & He, Y. Design of potent membrane fusion inhibitors against SARS-CoV-2, an emerging coronavirus with high fusogenic activity. J. Virol. 94 (2020).

[65] Gaurav, M., Sayrav, K., Kumar, A., Jha, D. K. & Yashvardhini, N. Genetic variations in the ORF7a protein of SARS-CoV-2 and its possible role in vaccine development. Biomed. Res. Ther. 8, 4497–4504 (2021).

[66] Gao, T. et al. Identification and functional analysis of the SARS-CoV-2 nucleocapsid protein. BMC Microbiol. 21, 58 (2021).

[67] Marino, S., Hogue, I. B., Ray, C. J. & Kirschner, D. E. A methodology for performing global uncertainty and sensitivity analysis in systems biology. J. Theor. Biol. 254, 178–196 (2008).

[68] R Development Core Team R: a language and environment for statistical computing. R Foundation for Statistical Computing (2011). https://www.r-project.org/

