## Supporting information for "Quantitative modeling of SARS-CoV-2 replication reveals phase-specific bottlenecks and antiviral targets"

**This file includes:**

Supporting text

Tables S1 to S3

Figures S1 to S10

SI references

### Supporting text

#### Protein normalization

Previous studies indicate that relative protein abundances measured by mass spectrometry are broadly consistent with absolute protein abundance distributions [1]. Based on this observation, and to reduce the number of fitted scaling parameters in the model, we assume that ratios of the MS measurements approximate ratios of absolute protein abundances across conditions. We utilized the western blot data to normalize the measurements of two MS datasets. The first dataset includes time kinetics (TK) 1–3, and the second includes TK 4 and 5. As these datasets are normalized separately, they are not directly comparable. Therefore, we normalized both datasets to N protein abundances at the 22 hours timepoint, which showed the lowest variance across all replicates within the plateau phase, and subsequently calculated the mean of both datasets. We then used the western blot nucleocapsid data at 22 hours to convert the intensities to absolute counts. Since the nucleocapsid western blot measurements are more accurate than the nucleocapsid MS measurements, we replaced the MS data with the western blot data for the N protein. The early measurements of the time series were removed for the measurement of the NSP proteins, as these depend solely on the peptides of NSP3 and thus cause a strong bias in the normalization of the MS values. The standard deviation was calculated and normalized separately for each MS dataset. When calculating the mean from the two datasets, we computed the weighted standard deviation of the respective individual standard deviations:

$$\sigma_{combined} = \sqrt{\frac{1}{n+m-1}((n-1)\sigma_1^2 + (m-1)\sigma_2^2 + \frac{mn}{m+n}(\mu_1 - \mu_2)^2)}.$$

$\sigma_i$ and $\mu_i$ represent the standard deviation and the mean of the datasets, respectively, while $n = 3$ and $m = 2$ indicate the respective number of data points in the datasets.

#### (sg)RNA length calculation

During transcription, it must also be taken into account that the different lengths of the individual sgRNAs significantly influence the replication dynamics. Every sgRNA starts with a 65bp leader sequence (ATTAAAGGTTTATACCTTCCCAGGTAA-CAAACCAACCAACTTTCGATCTCTTGTAGATCTGTTCT) followed by a 10bp transcription-regulatory-side-left (TRS-L) sequence (CTAAACGAAC) before each gen. We determined the start positions of each mRNA by the genome annotation file of the original SARS-COV-2 Wuhan strain (NCBI GenBank assembly GCA_009858895.3). The blue subsequence is used by Periscope [2] to determine the (sg)RNA count within our sequencing experiments.

#### Supplementary tables

**Table S1:** Total numbers of viral components predicted by the best-fit model, including non-packaged (+) gRNA strands, structural proteins, and both intracellular and extracellular infectious virus particles.

| Parameter | 6 hpi | 12 hpi | 24 hpi |
|---|---|---|---|
| (+)gRNA | 3145.04 | 28,714.11 | 30,698.36 |
| Intracellular virus | 111.92 | 1499 | 3665.70 |
| Extracellular virus | 2.56 | 212.30 | 1655.30 |
| N protein | 579,479.76 | 8,092,722.00 | 22,071,924.60 |
| S protein | 162,079.76 | 2,248,222.00 | 6,104,924.60 |
| M protein | 225,281.73 | 3,007,001.97 | 7,353,400.10 |
| E protein | 11,828.66 | 148,474.00 | 334,510.60 |

**Table S2:** Error between model predictions for montelukast's different modes of action and experimental data was quantified using the root mean square error (RMSE) and mean absolute error (MAE), calculated in $\log_{10}$-transformed space (Supplementary Fig. S4). **Bold** values indicate lowest error for each metric.

| Metric | NSP | Intracellular | Extracellular |
|---|---|---|---|
| RMSE ($\log_{10}$) | NSP1 | 1.35 | 0.893 |
| MAE ($\log_{10}$) | NSP1 | 0.868 | 0.607 |
| RMSE ($\log_{10}$) | NSP5 | **0.422** | **0.401** |
| MAE ($\log_{10}$) | NSP5 | **0.329** | **0.345** |

**Table S3:** Genome start positions and the corresponding (sg)RNA lengths based on the TRS sequence and the leader sequence.

| Gene | Parameter | Start position mRNA | RNA length |
|---|---|---|---|
| S | $l_S$ | 21563 | 8384 |
| ORF3a | $l_{3a}$ | 25393 | 4554 |
| E | $l_E$ | 26245 | 3702 |
| M | $l_M$ | 26523 | 3424 |
| ORF6 | $l_6$ | 27202 | 2745 |
| ORF7a | $l_{7a}$ | 27394 | 2553 |
| ORF8 | $l_8$ | 27894 | 2053 |
| N | $l_N$ | 28274 | 1673 |
| Genome | $l_G$ | 0 | 29903 |

#### Supplementary figures

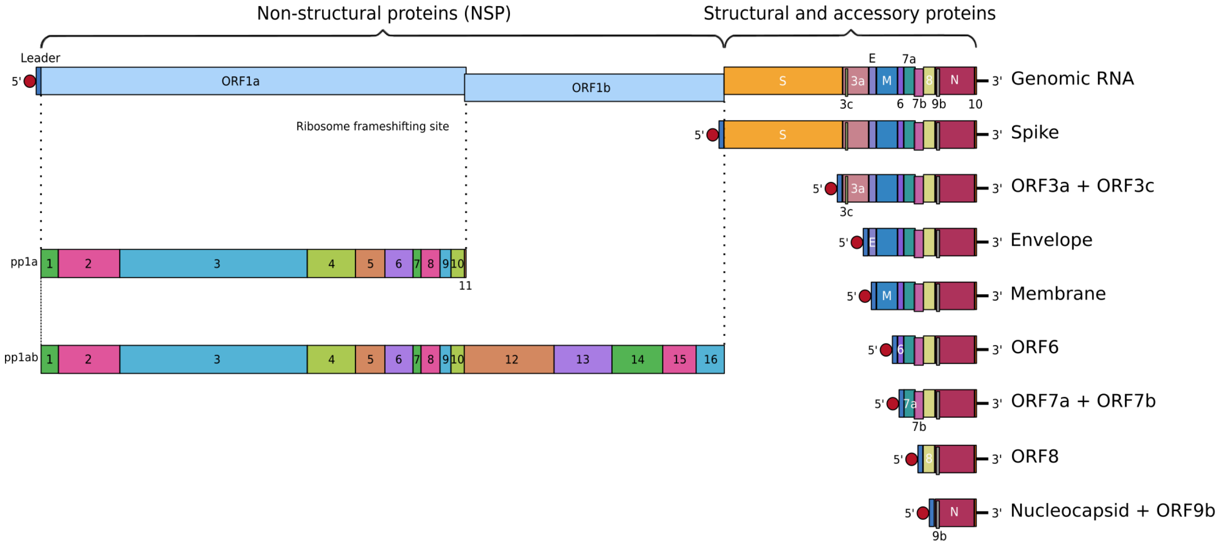


**Figure S1: SARS-CoV-2 genome organization.**
Schematic overview of the SARS-CoV-2 genome, including annotated subgenomic RNAs (sgRNAs) and the corresponding viral proteins expressed during infection. SARS-CoV-2 encodes four structural proteins: nucleocapsid (N), spike (S), membrane (M), and envelope (E); seven accessory proteins (ORF3a, ORF3c, ORF6, ORF7a, ORF7b, ORF8, and ORF9b); and two polyproteins (pp1a and pp1ab) that are further processed into 16 non-structural proteins (NSP1 through NSP16).

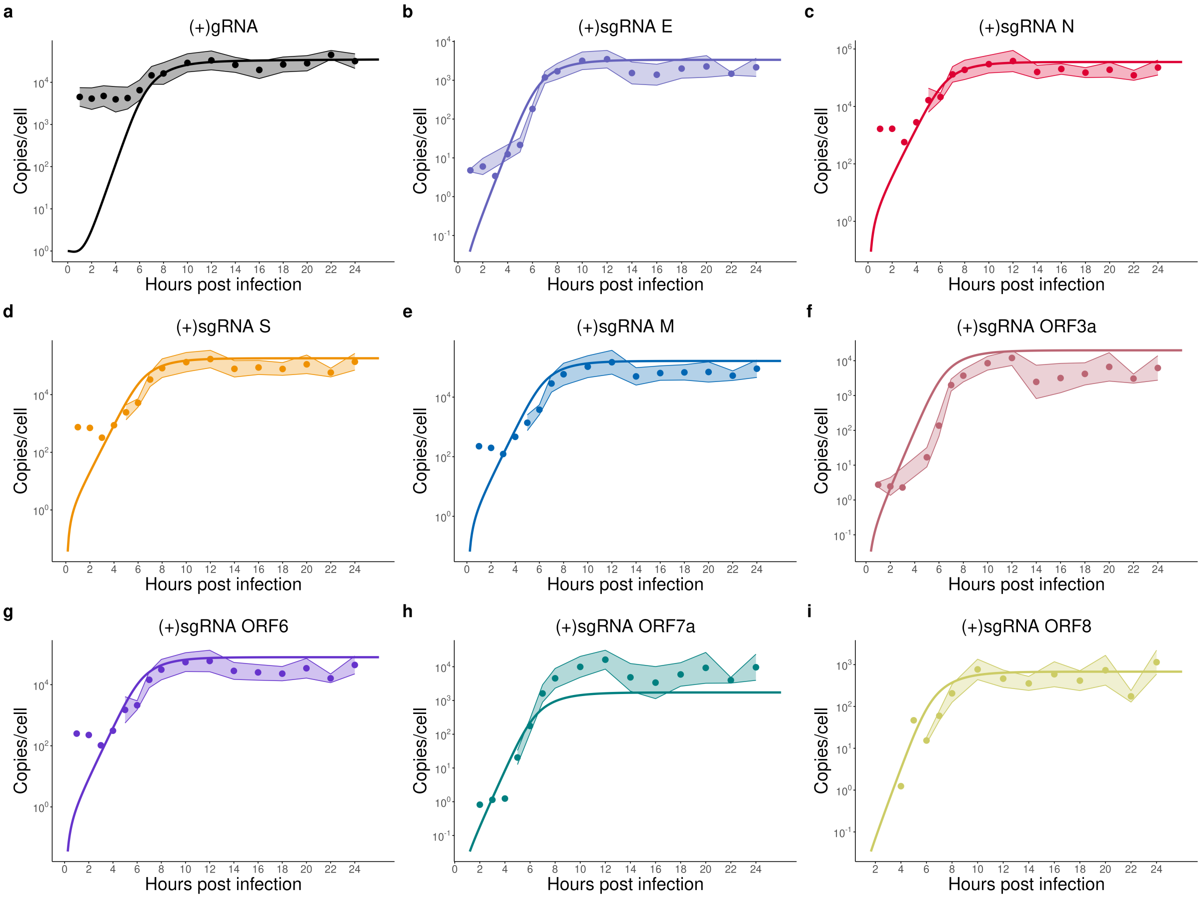


**Figure S2: Viral RNA species dynamics and model fits.**
Time-resolved RNA measurements and corresponding model fits are shown. **a** Genomic RNA (gRNA), quantified by RT-qPCR. **b–e** Subgenomic RNAs (sgRNAs) encoding structural proteins E, N, S, and M. **f–i** sgRNAs encoding accessory proteins ORF3a, ORF6, ORF7a, and ORF8 were detected by MinION sequencing and normalized to RT-qPCR measurements of E RNA (b). n=5. Shaded ribbons indicate 95% confidence intervals of the data.

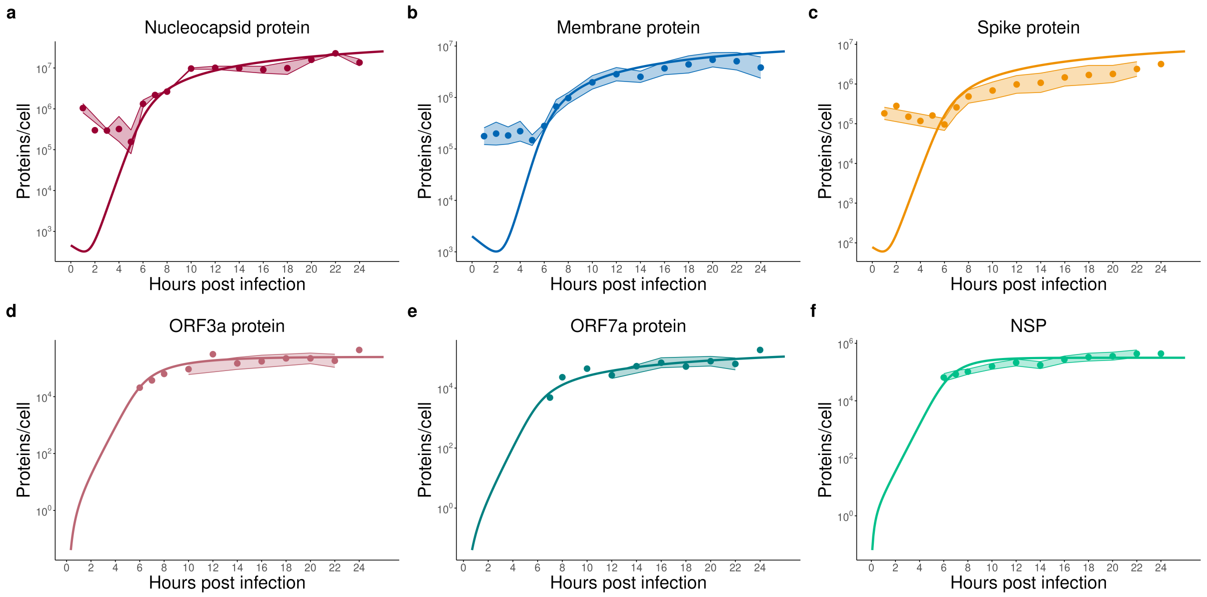


**Figure S3: Viral protein dynamics and model fits.**
Time-resolved protein abundance measurements and corresponding model fits are shown. **a**–**c** Viral structural proteins, including N, M, and S. **d**–**f** Non-structural proteins ORF3a, ORF7a, and NSP. Proteins such as E, ORF6, ORF8 were not quantified due to detection limits of the LC-MS/MS method. Shaded ribbons indicate 95% confidence intervals of the data. n=5.

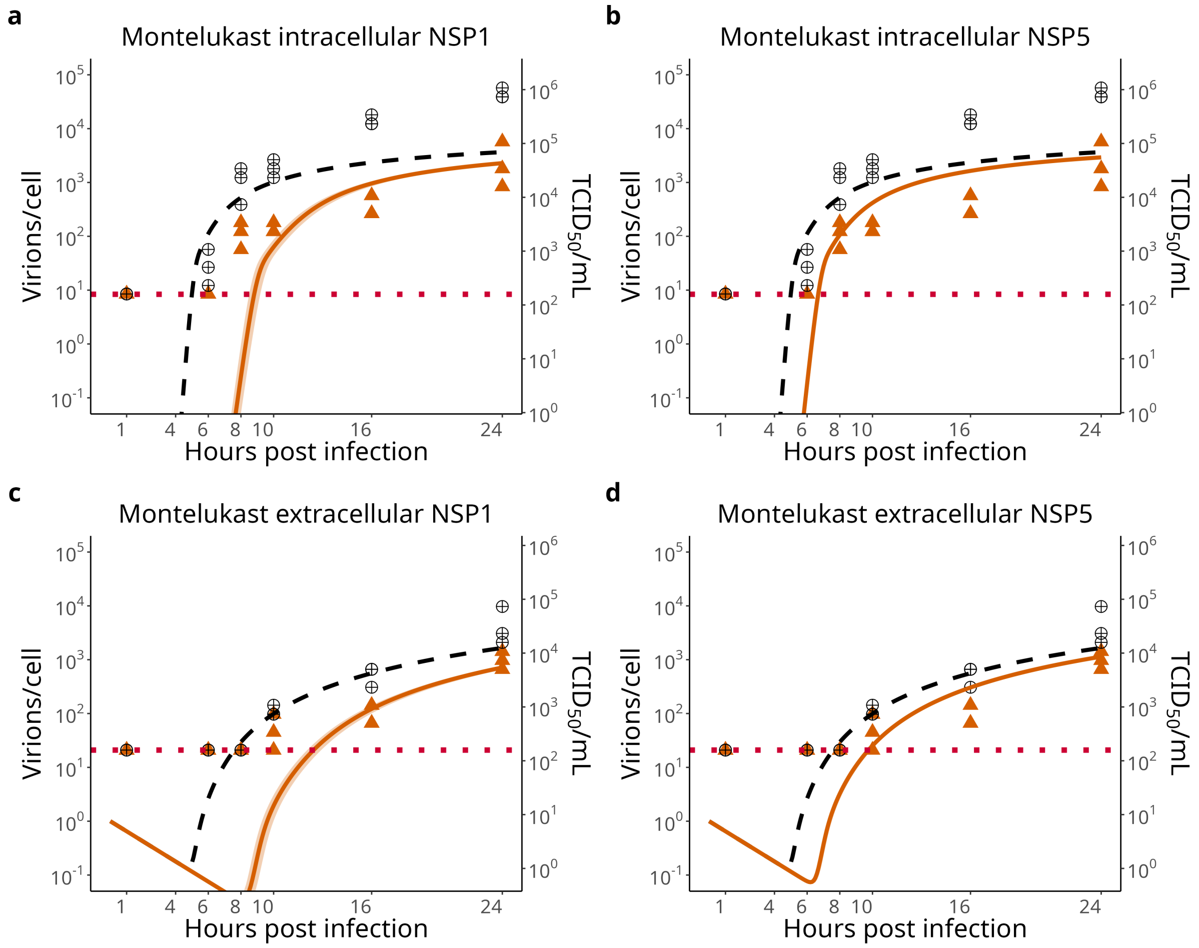


**Figure S4: Model comparison of montelukast targeting NSP1 and NSP5.** Simulated and experimental intracellular and extracellular virus titers under montelukast treatment are shown for two hypothesized targets. **a**, **c** Montelukast targeting NSP1. **b**, **d** Montelukast targeting NSP5. Both targeting scenarios reproduce the overall experimental trends, indicating that either mechanism can qualitatively explain the observed data.

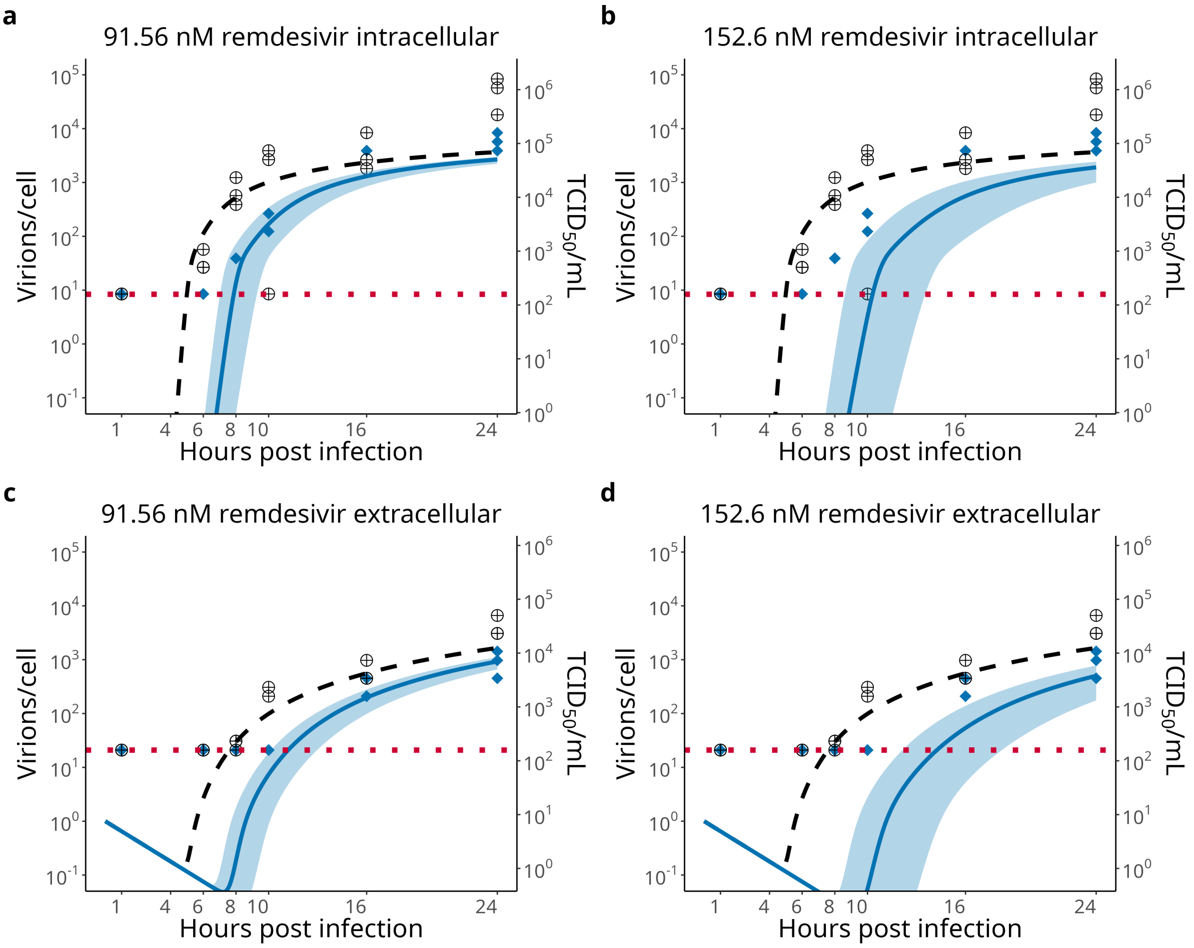


**Figure S5: Remdesivir treatment with reduced effective drug concentration.** Model simulations of remdesivir-mediated RdRp inhibition, implemented via the parameter $k_{RdRp}$, are shown for two effective inhibition levels. **a**, **c** 37.5% inhibition (91.56 nM). **b**, **d** 50% inhibition (152.5 nM). Simulations assuming 50% inhibition predict a stronger antiviral effect than observed experimentally, particularly at early time points.

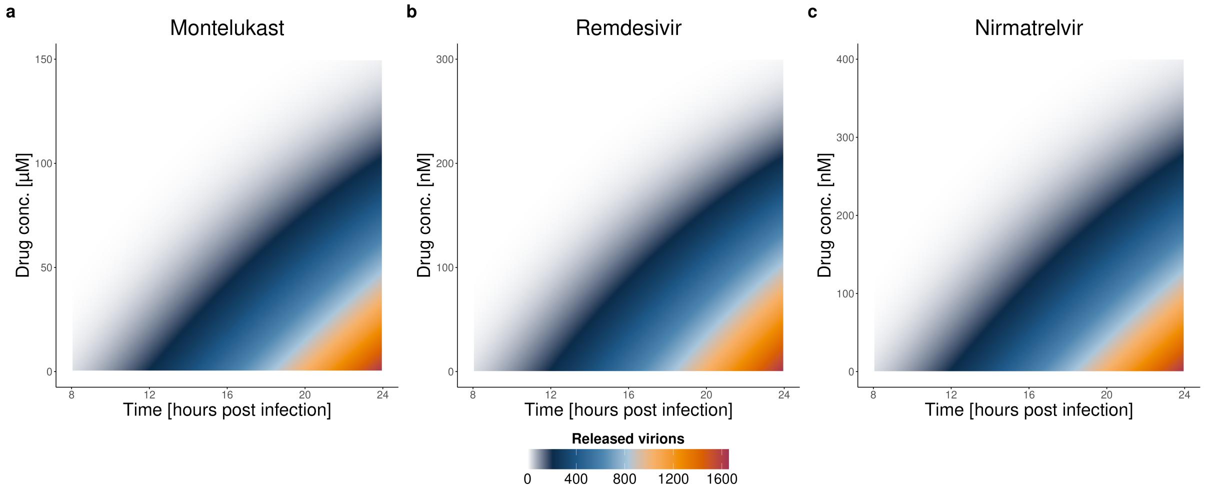


**Figure S6: Time-dependent effects of different antivirals.**
Modeled data illustrating time-dependent effects of different antiviral treatments with **a** montelukast, **b** remdesivir, and **c** nirmatrelvir on the release of virions from infected cells. The y-axes represent drug concentration, the x-axes represent time post-infection, and the color gradient indicates the relative amount of released virions (white = none, dark red = more virions). Treatment delays the onset of viral release, with the extent of delay increasing with drug concentration, but does not completely eliminate viral production.

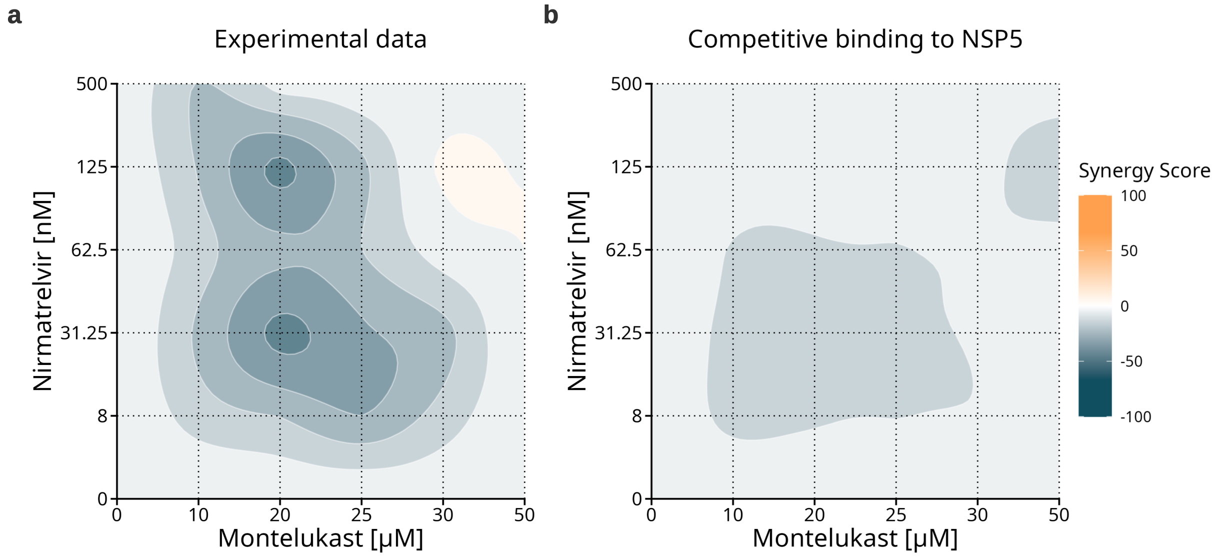


**Figure S7: Loewe synergy analysis of nirmatrelvir and montelukast combination treatment.** Loewe synergy scores were calculated to assess drug–drug interactions under the assumption of a shared target and mechanism of action. **a** Experimental data. **b** Model simulations. For the simulations, we assumed antagonistic interactions of the two drugs in which the combined drug effect was represented by the stronger inhibitory effect of either compound on the shared target.

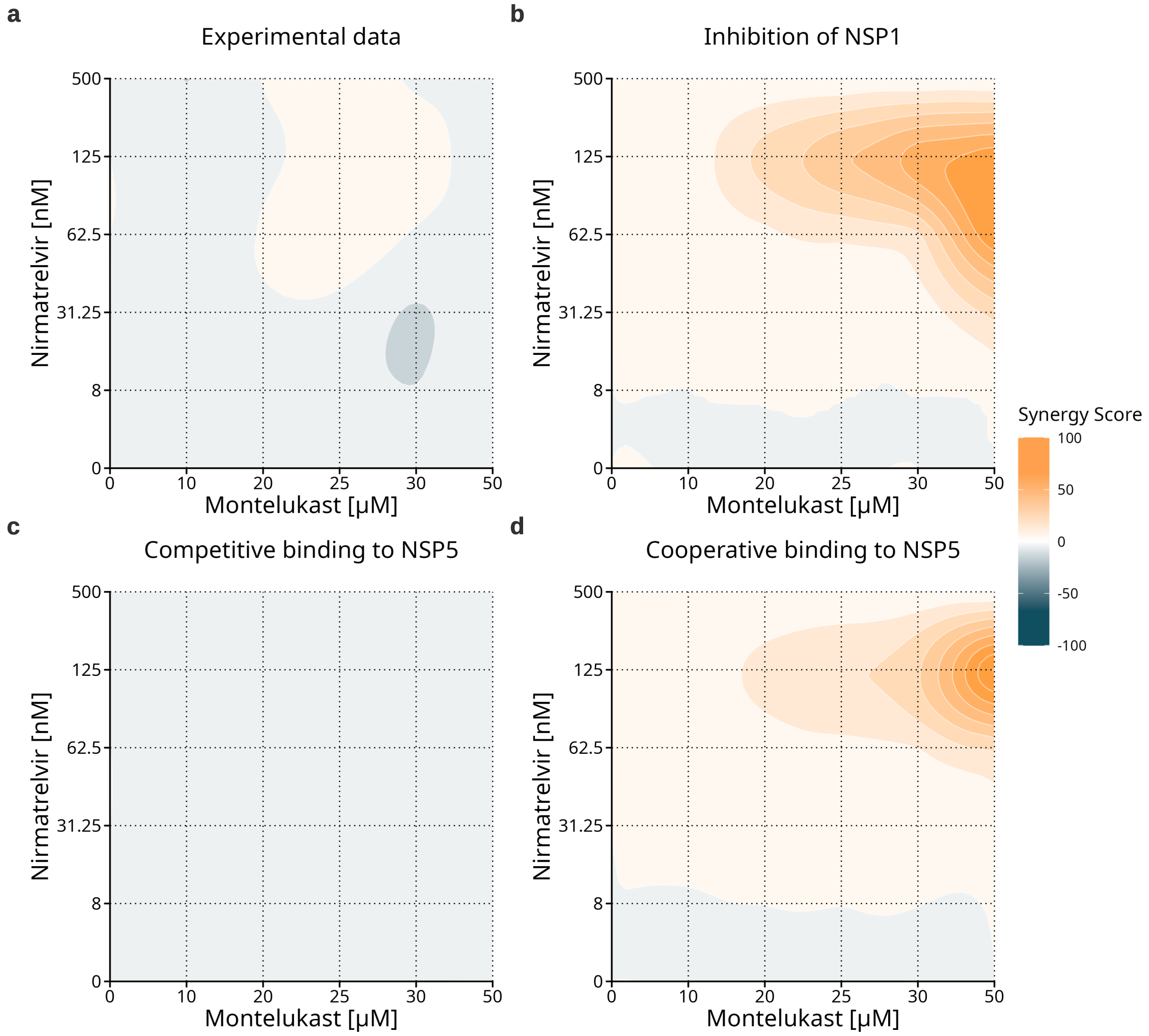


**Figure S8: ZIP synergy analysis of nirmatrelvir and montelukast under different target assumptions.**

ZIP synergy scores were calculated for combination treatment with nirmatrelvir and montelukast under different mechanistic scenarios. Nirmatrelvir was assumed to target NSP5 in all cases. **a** Experimental data. **b** Montelukast targeting NSP1 via inhibition of $k_1$. **c** Montelukast targeting NSP5 with an antagonistic interaction, where only the stronger inhibitory effect of either drug is applied. **d** Montelukast targeting NSP5 with a synergistic interaction, where both inhibition functions (as described in the M&M paragraph) are applied to $p_{NSP}$. Model simulations indicate that the antagonistic interaction scenario **c** provides the closest agreement with the experimental data, supporting NSP5 as the primary target of montelukast and suggesting competitive or interfering drug effects.

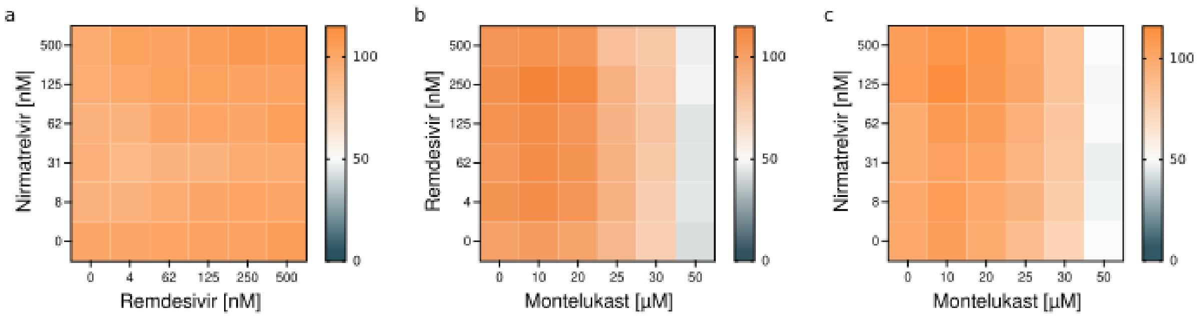


**Figure S9: Cytotoxic effects of antiviral combination treatments assessed by MTT assay.** Heat maps show the cytotoxic effects of antiviral combination treatments at indicated concentrations, as determined by MTT assays (24 h post-treatment). **a** Nirmatrelvir and remdesivir, **b** remdesivir and montelukast, and **c** nirmatrelvir and montelukast. Colors represent the relative viability (0-100%) for each treatment combination (normalized to the DMSO control treatment). n=3.

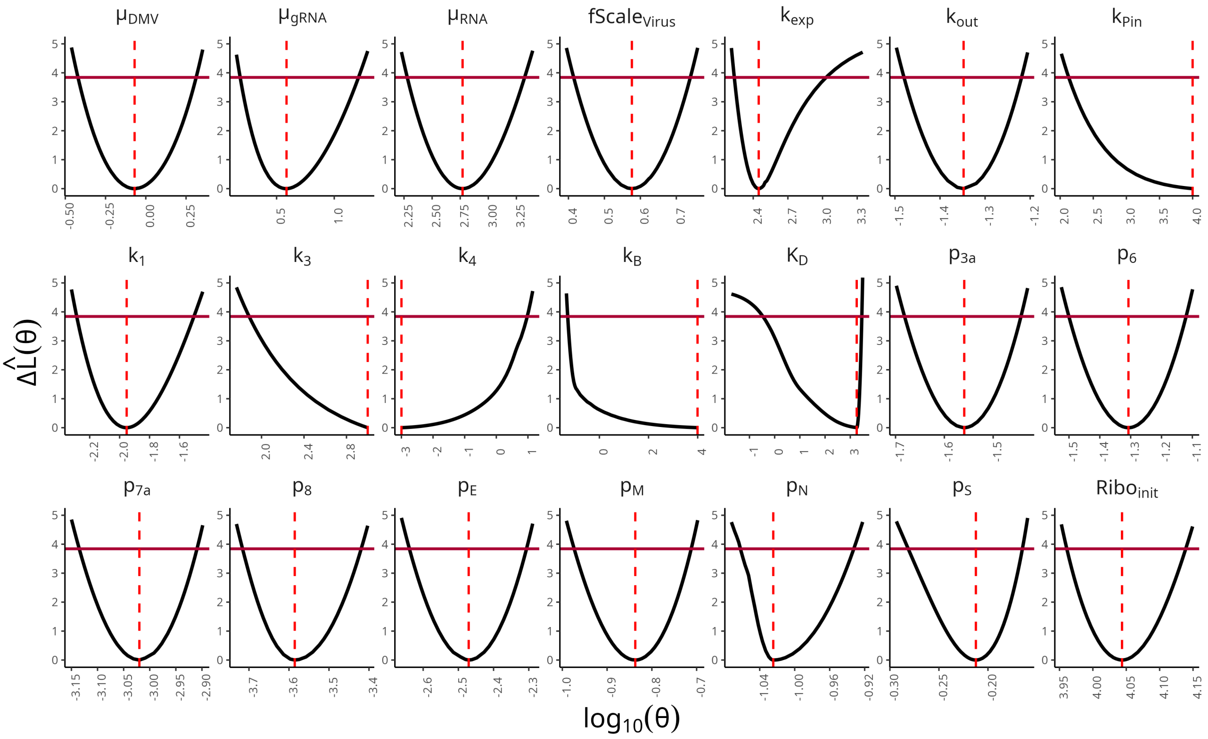


**Figure S10: Profile likelihood analysis of model parameters.**
Profile likelihoods for all estimated model parameters $\theta$ are shown as black curves. The red lines indicate the statistical 95% threshold for each parameter. Intersections of the profile likelihood curves with this threshold define the 95% confidence intervals, which are listed in Table 1. Parameters with finite confidence intervals are considered practically identifiable. The dashed red lines mark the optimal parameter value, also detailed in Table 1. The x-axis displays parameter values on a $\log_{10}$ scale, while the y-axis shows the change in negative log-likelihood $\hat{L}(\theta)$.

### References

[1] Rais, Y., Fu, Z. & Drabovich, A. P. Mass spectrometry-based proteomics in basic and translational research of SARS-CoV-2 coronavirus and its emerging mutants. *Clin. Proteomics* **18**, 19 (2021).

[2] Parker, M. D. et al. Subgenomic RNA identification in SARS-CoV-2 genomic sequencing data. *Genome Res.* **31**, 645–658 (2021).
